# Chromatin regulatory dynamics of early human small intestinal development using a directed differentiation model

**DOI:** 10.1101/2019.12.18.881219

**Authors:** Yu-Han Hung, Sha Huang, Michael K. Dame, Qianhui Yu, Qing Cissy Yu, Yi Arial Zeng, J. Gray Camp, Jason R. Spence, Praveen Sethupathy

## Abstract

**Background:** The establishment of the small intestinal (SI) lineage during human embryogenesis is critical for the proper development of neonatal gut functions, including nutrient absorption and immune defense. The chromatin dynamics and regulatory networks that drive human SI lineage formation and regional patterning are essentially unknown. To fill this knowledge void, we apply a cutting-edge genomic technology to a state-of-the-art human model of early SI development. Specifically, we leverage chromatin run-on sequencing (ChRO-seq) to define the landscape of active promoters, enhancers, super enhancers, and gene bodies across distinct stages of directed differentiation of human pluripotent stem cells (hPSCs) into SI spheroids with regional specification.

**Results:** Through comprehensive ChRO-seq analysis we identify candidate stage-specific chromatin activity states, novel markers, and enhancer hotspots during the directed differentiation process. Moreover, we propose a detailed transcriptional network associated with SI lineage formation or initial regional patterning. Among our findings is a unique pattern of enhancer activity and transcription at HOX gene loci that is previously undescribed. Analysis of single cell RNA-seq data from human fetal SI at early developmental time points shed further light on the unique HOX gene temporal dynamics that underlies SI regional patterning.

**Conclusions:** Overall, the results lead to a new proposed working model for the regulatory underpinnings of human SI lineage formation and regional patterning, thereby adding a novel dimension to the literature that has thus far relied almost exclusively on non-human models.

## Introduction

The embryonic development of the small intestine (SI) is critical for a fetus to thrive and grow. The genetic programming of SI cell identity and regional specification during early development is fundamental to ensure proper SI morphogenesis and maturation with specialized functions, including nutrient digestion and absorption, energy balance, and pathogen defense. Several seminal studies have identified important molecular regulators associated with gut development [1-5], and more recent studies have leveraged advanced genomic technologies (e.g., single cell RNA-seq and bulk ATAC-seq) to provide insights at a more granular level into gut development [6-11]; however, this work relies almost exclusively on animal models. Studies of human SI development have been few [7, 12, 13], due in large part to limited access to primary human fetal tissues. Initial human SI lineage formation from the endodermal germ layer, which occurs much prior to the developmental time points at which human fetal tissues are generally available, is essentially uncharacterized. In this study, we sought to fill this important knowledge gap by profiling for the first time the chromatin regulatory dynamics and transcriptional programs that are associated with human SI lineage formation and initial regional patterning.

Robust temporal and spatial regulation of gene expression is fundamental to all biological processes including development [14, 15]. Transcriptional programs are precisely controlled by promoters and distal *cis*-regulatory regions known as enhancers. Enhancers harbor binding sites for transcription factors (TFs), activate long-range gene transcription, and are often cell-type specific [16, 17]. RNA polymerases are known to be recruited to active enhancers, generating divergent short transcripts (also known as enhancer RNA or eRNAs) [18, 19]. Recently, an approach called chromatin run-on sequencing (ChRO-seq) [20] was developed for genome-wide identification of promoters, active enhancers, and actively transcribed gene bodies in a single assay. ChRO-seq represents the newest generation of nascent RNA sequencing technologies, and overcomes several limitations of previous versions including global run-on sequencing (GRO-seq) [21] and precision run-on sequencing (PRO-seq) [22]. ChRO-seq was very recently successfully applied to archived solid tumor tissues [20, 23].

Advances in the directed differentiation of human pluripotent stem cells (hPSCs), including human embryonic stem cells (hESCs), provide a powerful strategy for studying the early developmental events of human SI that would be nearly impossible otherwise [24, 25]. The multi-step process of generating human “gut-in-a-dish” from hPSCs recapitulates many aspects of *in vivo* SI development [26-28], including induction of the endodermal fate, formation of SI lineage (SI spheroids) with regional identities, and maturation into 3-dimensional human intestinal organoids (HIOs) that exhibit molecular, structural, and functional features similar to those of the human fetal SI [13, 29]. While the protocol for directed differentiation of hPSCs into SI spheroids has been established, the molecular mechanisms governing this process remains minimally characterized. To reveal temporal dynamics of the chromatin state during early human SI development and regional patterning, we performed ChRO-seq across the following stages: hPSC, definitive endoderm (DE), duodenal (proximal SI) spheroid, and ileal (distal SI) spheroid. This study provides the first-ever view of the changing chromatin regulatory landscape, defines stage-specific enhancer hotspots, and identifies key TF networks that underlie the acquisition of SI identity and the initiation of ileal regional patterning. Moreover, we uncover previously undescribed dynamics at HOX gene loci associated with these events. Additional analysis of single cell RNA-seq (scRNA-seq) of human fetal SI tissue underscores the unique HOX gene patterns, although these results are subject to the caveat that these tissues reflect developmental time points much later than what we have the opportunity to analyze in the directed differentiation model. Overall, this study offers an unprecedented resource that serves as a springboard from which the research community can develop and test targeted hypotheses about key regulatory hotspots and molecular drivers of early events of human SI development.

## Results

### ChRO-seq reveals temporal dynamics of nascent transcription at gene loci in the directed differentiation model of human developing SI

The directed differentiation of hPSCs (here we used H9 hESCs) into SI spheroids with regional specification was carried out as previously described [26, 28] **(Figure 1A)**. The patterned SI spheroids represent the primitive form of the SI comprised of mainly stem/progenitor cells [30], which can give rise to different mature epithelial cell types after prolonged organoid culture, or following transplantation into a murine host [31]. In this study, we included the four stages of this model: hESC, DE, duodenum (Duo) spheroid, and ileal (Ile) spheroid **(Figure 1A)**. We carried out ChRO-seq in order to characterize the dynamics of genome-wide nascent transcriptional activity toward the goal of defining the changing landscape of promoters, enhancers, and active gene bodies across the following sequential events: DE fate induction (from hESC to DE), SI lineage formation (from DE to Duo spheroid), and initial ileal regional patterning (from Duo to Ile spheroid) **(Figure 1A)**.

**Figure 1.**
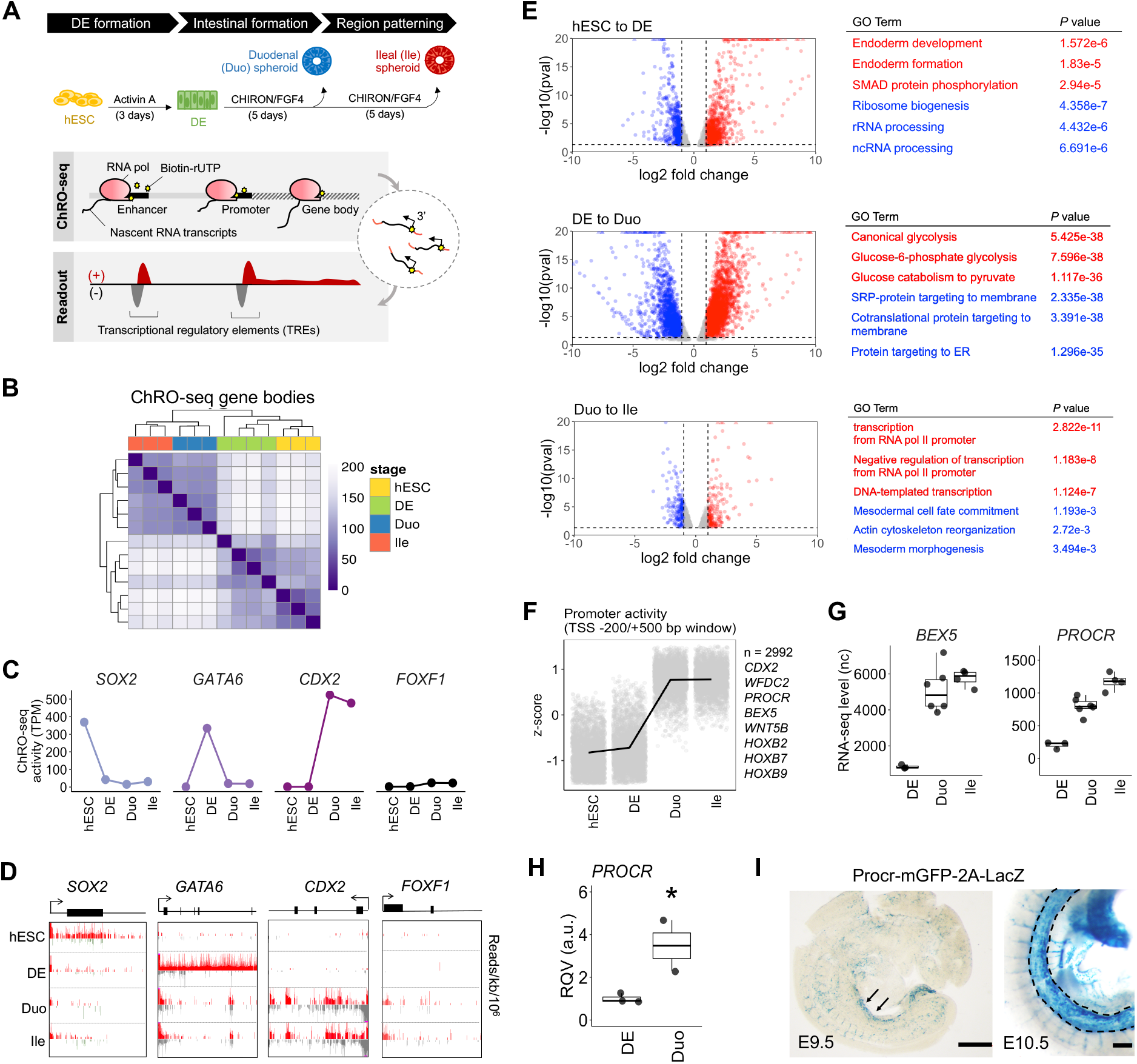
ChRO-seq defines dynamics of nascent transcription at promoters and gene loci during directed differentiation from human pluripotent stem cells (hPSCs) to small intestinal (SI) spheroids with regional specification. (A) Schematic diagram of the distinct stages of directed differentiation and the genome-scale approach of ChRO-seq. (B) Hierarchical clustering analysis of gene transcription profiles across all stages of the model. (C) Transcriptional activity (ChRO-seq signal) of the genes *SOX2, GATA6, CDX2*, and *FOXF1* across all stages of the model. (D) Normalized ChRO-seq signal at the gene bodies of *SOX2, GATA6, CDX2* and *FOXF1* across all stages of the model. Scale (+/- 25) is fixed across stages. (E) Volcano plots showing differentially transcribed genes in the indicated comparisons (average TPM > 25, log_2_ fold change of transcription > 1, padj < 0.2 and p < 0.05 by Wald test; DESeq2). Results of pathway enrichment analyses of up-transcribed (red) and down-transcribed (blue) genes in the indicated comparisons (GO Biological Process 2018) are also shown. (F) Identification of protein-coding genes that have high promoter activity in SI spheroids (n = 2,992) through application of the likelihood ratio test across all stages. (G) Expression levels (RNA-seq) of *BEX5* and *PROCR* in stages of DE and SI spheroids. (H) RT-qPCR of *PROCR* in an independent batch of samples (DE = 3; Duo spheroid = 2). *P < 0.05 by one-tailed t test (validation of sequencing data). (I) X-gal staining for β-galactosidase activity of Procr-mGFP-2A-LacZ mouse reporter line showing a robust *Procr* expression in midgut at E9.5 (left; arrowhead) and E10.5 (right; dashed outline). Scale bar = 500 μm. ChRO-seq study: hESC, n = 3; DE, n = 4; Duo spheroid (Duo), n = 3; Ile spheroid (Ile), n = 3. RNA-seq study: hESC, n = 2; DE, n = 3; Duo, n = 6; Ile, n = 4. TPM, transcripts per million. nc, normalized counts. RQV, relative quantitative value.

To assess nascent transcription of genes, we first analyzed ChRO-seq signal within the body of annotated genes. ChRO-seq data in the first 500 bp downstream of the transcription start site, or TSS, was excluded to remove signal contributed by RNA polymerase pausing. Hierarchical clustering analysis of the gene transcriptional profiles demonstrates clean stratification of the different stages **(Figure 1B)**. The signal at gene bodies encoding TFs that are well-established markers of particular stages (*SOX2, GATA6*, and *CDX2*) indicate the expected specificity **(Figure 1C-D)**. Also, the very minimal expression of the mesenchymal marker *FOXF1* **(Figure 1C-D)**, as well as markers of other organ lineages **(Supplementary Figure 3B)**, demonstrate that the Duo and Ile spheroids are indeed dominated by early-stage SI epithelial cells. Next we sought to identify from the ChRO-seq data differentially transcribed genes during each stage transition of this model (log_2_ fold change > 1, average TPM > 25, padj < 0.2, p < 0.05 by Wald test; DESeq2) **(Figure 1E)**. The analyses identified 1886 differentially transcribed genes (1328 up; 558 down) during DE formation (DE vs. hESC), 4024 differentially transcribed genes (2546 up; 1478 down) during SI lineage formation (Duo spheroids vs. DE) and 362 differentially transcribed genes (182 up; 180 down) during ileal patterning event (Ile vs. Duo spheroids) **(Figure 1E)**. We also performed RNA-seq on the same stages to define differentially expressed genes and the Gene Ontology (GO) term enrichment analysis between differentially transcribed and differentially expressed genes are in general agreement **(Supplementary Figure 2)**.

The same ChRO-seq data also affords the unique opportunity to evaluate promoter activity. Therefore, we next developed an approach to analyze promoter dynamics across all of the stages **(Supplementary Figure 1C)**. We identified 9550 genes, the majority of which are protein-coding genes (n = 7651), which exhibit significantly changing patterns of promoter activity across different stages (padj < 0.05 by likelihood ratio test; DESeq2) **(Supplementary Figure 1C)**. Notably, a subset of these protein-coding genes has low promoter activity in both hESC and DE but high promoter activity in both Duo and Ile spheroids (n=2992). These include *BEX5* and *PROCR* **(Figure 1F)**, which were recently observed to have robust expression in primary human fetal SI at early developmental time points [7, 12]. Analysis of the matched RNA-seq data, which showed a dramatic rise in *BEX5* and *PROCR* mRNA levels during formation of SI spheroids from DE **(Figure 1G)**, confirm the findings from the promoter analysis. We also validated the upregulation of *PROCR* in SI spheroids by RT-qPCR with an independent batch of samples **(Figure 1H)** and demonstrated by *in situ* hybridization staining a robust expression of *Procr* in the midgut region of *Procr*-mGFP-2A-LacZ mice [32] at E9.5 and E10.5 (**Figure 2I**), indicating the conserved early presence of *PROCR* in the developing SI. Overall, these findings demonstrate that hPSC-derived SI spheroids recapitulate features of early developing SI and therefore represent the state-of-the-art model for further study of human SI lineage formation.

**Figure 2.**
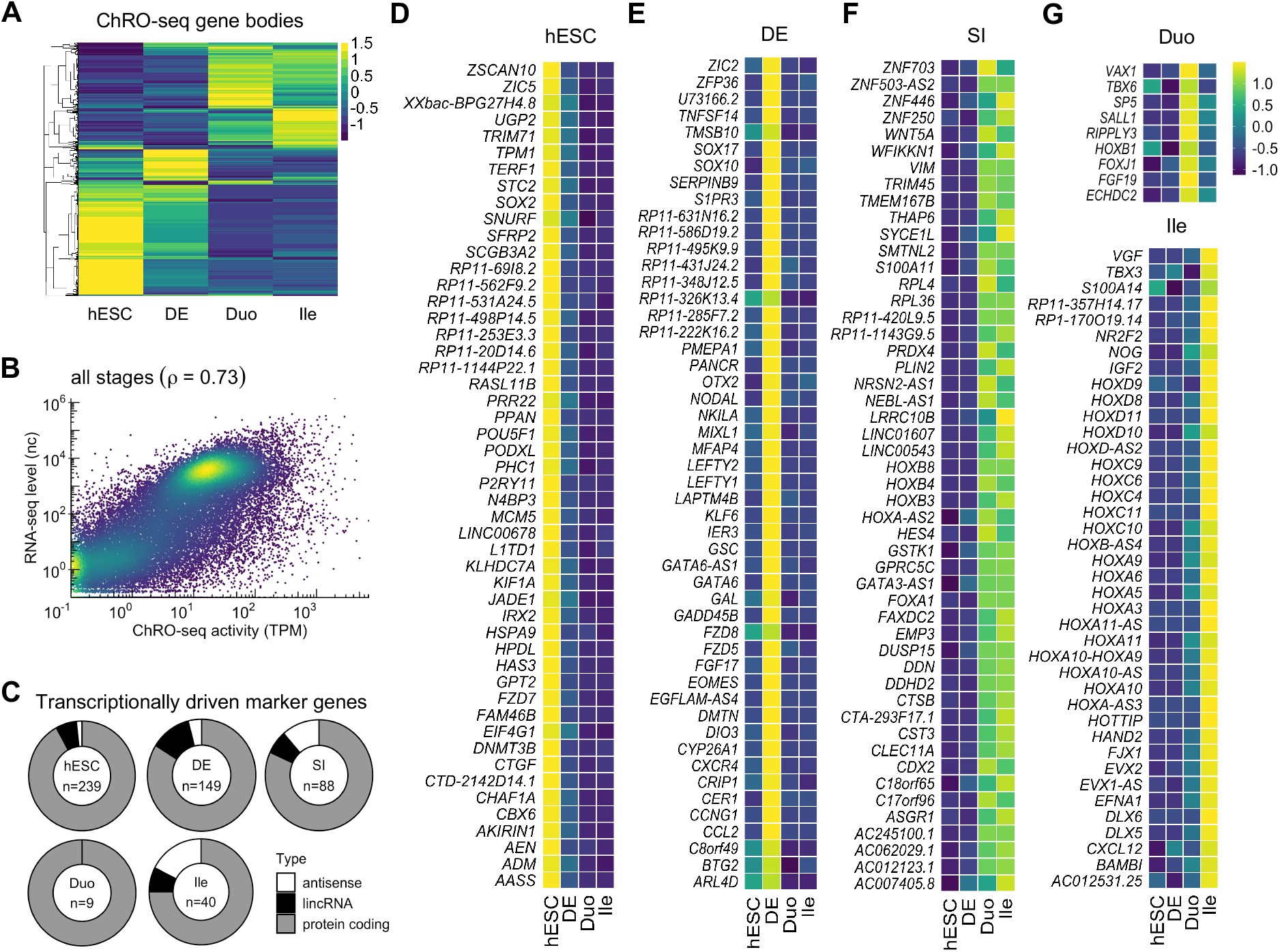
Identification of transcriptional markers of specific stages in the directed differentiation model of human developing SI. (A) Hierarchical clustering analysis of transcribed genes (TPM > 50 at least in one stage) (z-score). (B) Genome-wide correlation of transcribed levels (ChRO-seq) and expressed levels (RNA-seq) of genes. Transcription and expression levels of genes were averaged across all stages. No transcription and expression thresholds were used (see also Supplementary **Figure 3B-E**). (C) Identification of markers that label specific stages at the levels of both transcription and steady-state expression. The total number of markers (center of the donut plot) and the proportion of marker types are shown for each stage. (D-G) Marker genes of hESC (D), DE (E), SI spheroids regardless of regional identity (F), Duo and Ile (G). In panel (D-F), only the top 50 most variable genes across stages are shown (see **Supplementary Data 2** for full list). Heatmaps denote variation across stages (z-score). ChRO-seq study: hESC, n = 3; DE, n = 4; Duo spheroid (Duo), n = 3; Ile spheroid (Ile), n = 3. RNA-seq study: hESC, n = 2; DE, n = 3; Duo, n = 6; Ile, n = 4.

### Identification of transcriptional markers of specific stages in the directed differentiation model of human developing SI

ChRO-seq signal at gene bodies (TPM > 50 in at least one stage) reveals the changing patterns of transcription across stages **(Figure 2A)**. We sought to identify transcriptional markers of each stage in this model. It is important to note that previously validated SI regional markers using human fetal duodenum and ileum ranging 14-19 weeks of gestation [28] are not sufficient to distinguish Duo from Ile spheroids, although they are sufficient to distinguish Duo from Ile HIOs (derived after culturing the spheroids in Matrigel for 28 days) **(Supplementary Figure 3A)**. This likely reflects the fact that spheroids, which are newly differentiated, represent the early gut lineage whereas the HIOs represent a more mature state. Here, by leveraging ChRO-seq signal at gene bodies as well as RNA-seq data, we were able to define gene sets that serve as early markers of SI lineage formation and ileal regional patterning at the levels of both transcription and steady-state expression.

We first demonstrated that ChRO-seq and RNA-seq signal for genes are reasonably well-correlated (Pearson correlation coefficient = 0.73) **(Figure 2B; Supplementary Figure 3B-E)**, indicating that nascent transcription profiles globally reflect steady state expression profiles in this model system (albeit not perfectly, as expected due to other layers of gene regulation such as those at the post-transcriptional level). To identify genes that label a specific stage at the levels of both transcription and steady-state expression, we developed a bioinformatic pipeline to integrate ChRO-seq and RNA-seq data (Methods) and performed this integrative analysis with genes that are actively transcribed (TPM > 50) in at least one stage. We identified genes that are significantly elevated according to both transcription and steady-state expression in each stage relative to all other stages: hESC (n=239), DE (n=149), SI spheroid irrespective of regional identity (n=88), Duo spheroid (n=9), and Ile spheroid (n=40) **(Figure 2C; Supplementary Data 2)**. The stage-specific genes we identified include well-established markers such as *POU5F1* and *SOX2* for hESCs **(Figure 2D)**; *NODAL, SOX17, EOMES* and *LEFTY2* for DE **(Figure 2E)**; and *CDX2* for SI spheroids irrespective of regional identity **(Figure 2F)**. We defined additional stage-specific markers, many of which are previously undescribed: *HES4, FOXA1, WFDC2*, and several *HOXB* family members for SI spheroids irrespective of regional identity; *SP5* and *SALL1* for Duo spheroids; and *FJX1, IGF2*, as well as several HOXA/C/D members for Ile spheroids **(Figure 2F)**. We also identified several long, non-coding RNA (lncRNA) and anti-sense transcript markers: *LINC00543* for SI spheroids irrespective of regional identity, and *EVX1-AS* for Ile spheroids **(Figure 2C-F)**. Notably, while many genes were defined as markers of SI spheroids irrespective of regional identity, the analysis identified much fewer genes as Duo-specific markers compared than Ile-specific markers **(Figure 2C)**, which may suggest that Ile regional specification is mainly driven by gain of additional driver genes rather than loss of genes contributing to Duo regional identity.

### Transcriptional regulatory element profiles reveal notable re-wiring of chromatin activity during directed differentiation

Active transcriptional regulatory elements (TREs), including promoters and enhancers, are identified in ChRO-seq data by the hallmark feature of short bi-directional transcription **(Figure 3A)**. To identify active TREs across the entire genome, we employed dREG [33], which was developed specifically for this purpose. Using this tool, we identified a total of 125,863 active TREs across all four stages included in this study **(Figure 3B)**. The length distribution of these active TREs is consistent with what has been reported previously [34] and the vast majority of the active TREs, as expected, are located in intergenic regions, introns, and annotated TSSs **(Figure 3C-D)**. To specifically evaluate whether the TRE landscape of SI spheroids is more similar to developing as opposed to mature SI in humans, we leveraged data on DNase-accessible regions in both fetal and adult human SI available from the Roadmap Epigenomics Project. We first defined DNase-accessible regions unique to fetal or adult tissue (Methods) and then intersected those with active TREs present in Duo and/or Ile spheroids. We found that 12,541 out of 37, 892 (∼33%) open chromatin loci specific to fetal human SI overlap with TREs present in Duo and/or Ile spheroids, whereas only 7,504 out of 45,242 (∼16%) open chromatin loci specific to adult human SI overlap with TREs present in Duo and/or Ile spheroids. To provide a specific example of a TRE defined in this model that captures an important regulatory region associated with human fetal SI, we turned to the very recently identified intestinal critical region (ICR). The ICR represents an important regulatory element of the new gene *PERCC1*, which exhibits transient expression in the SI during development and has been linked to congenital diarrheal disorders in humans [35]. We identified a TRE that is highly overlapping with the ICR and active only in SI spheroids, especially in Duo spheroids **(Figure 3F-G)**. Taken together, these observations demonstrate that TREs in hPSC-derived SI spheroids are able to capture important features of gene regulatory landscapes that are associated with the developing human SI.

**Figure 3.**
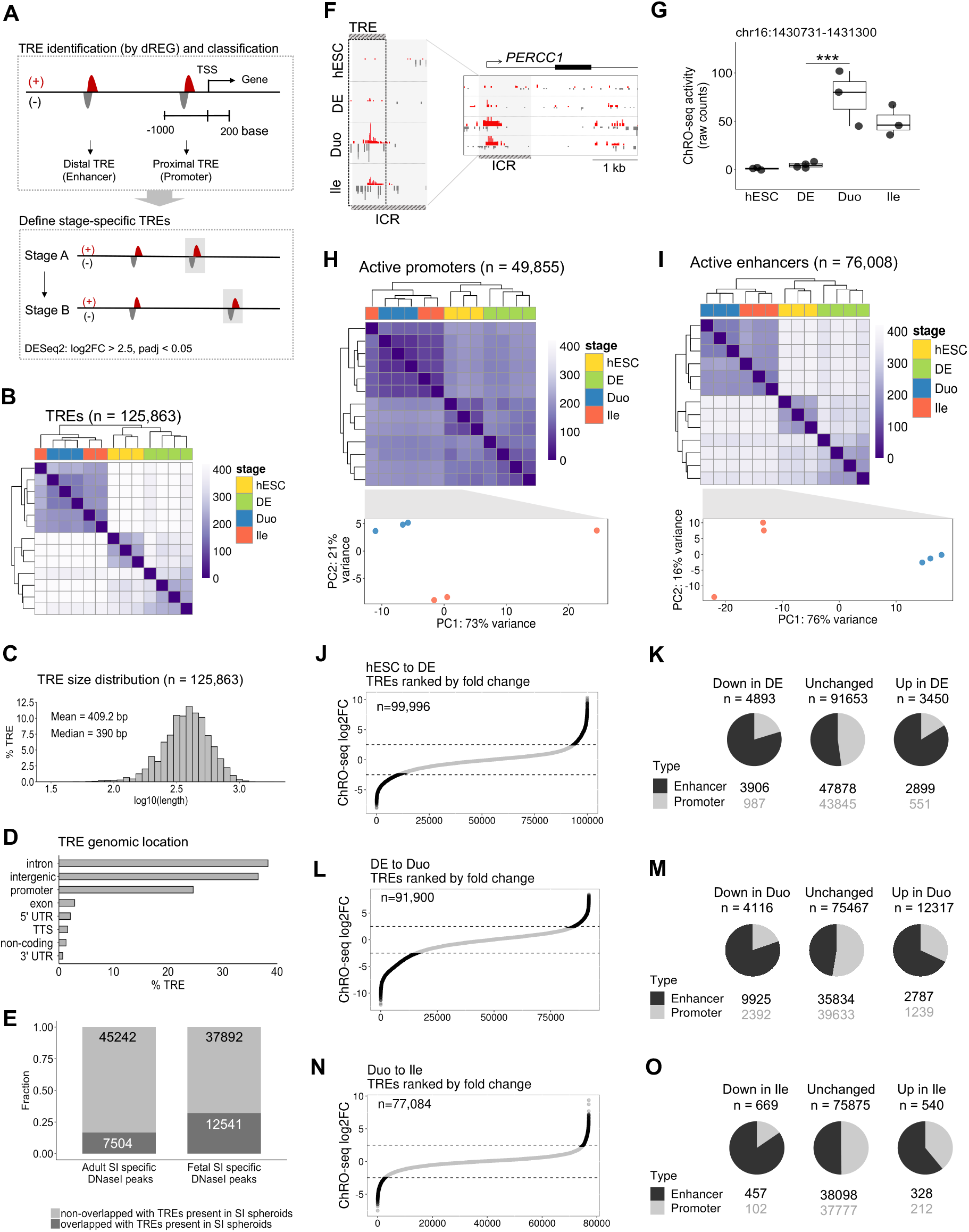
Transcriptional regulatory element profiles reveal notable re-wiring of chromatin activity in the directed differentiation model of human developing SI. (A) Schematic for identification and categorization of active TREs, followed by definition of TREs with altered activity during stage transitions of the model. (B) Hierarchical clustering analysis of profiles of all active TREs. (C) Size distribution of active TREs. (D) Genomic locations of active TREs (hg38 build). (E) Percentage of fetal and adult SI-specific open chromatin regions (defined according to DNaseI-seq data from the Roadmap Epigenomics Project) overlapping with active TREs present in Duo and/or Ile spheroids. (F) ChRO-seq signal around the Intestinal Critical Region (ICR) [35]. An active TRE identified in our study (chr16: 1430731-1431300; hg38) largely overlaps with the ICR and exhibits strong ChRO-seq signals in SI spheroids. Scales in right panel (+25/-25) and left panel (+166/-15; the maximum range) are fixed across different stages. (G) The transcriptional activity of the TRE associated with the ICR (chr16: 1430731-1431300; hg38) across all stages of the model (*** P < 0.001 by Wald test; DESeq2). (H) Hierarchical clustering (upper) and PCA (lower) analysis of promoter profiles in the indicated stage. (I) (H) Hierarchical clustering (upper) and PCA (lower) analysis of enhancer profiles in the indicated stage. (J) Active TREs present in hESC and DE (n = 99,996) and ranked based on the log2 fold change in activity during the transition to DE. (K) Numbers of promoters and enhancers that are unchanged, gained (up), or lost (down) during the transition from hESC to DE. (L) Active TREs present in DE and Duo spheroid (n = 91,900) and ranked based on the log2 fold change in activity during Duo spheroid formation. (M) Numbers of promoters and enhancers that are unchanged, gained (up), or lost (down) during the transition from DE to Duo spheroid. (N) Active TREs present in Duo and Ile spheroid (n = 77,084) and ranked based on the log2 fold change in activity during Ile spheroid formation. (O) Numbers of promoters and enhancers that are unchanged, gained (up), or lost (down) in Ile compared to Duo spheroids. hESC, n = 3; DE, n = 4; Duo spheroid (Duo), n = 3; Ile spheroid (Ile), n = 3.

We next categorized the identified active TREs into 49,855 proximal and 76,008 distal TREs, which from here on in we refer to as promoters and enhancers, respectively **(Figure 3A)**. Unsupervised hierarchical clustering analyses reveal that enhancer profiles stratify different stages more accurately and clearly than promoters **(Figure 3H-I)**, consistent with the notion that enhancer signature is the most cell-type specific [14, 16]. To further elucidate the dynamics of active TRE profiles across the directed differentiation process, we identified active TREs that are unchanged, gained (enhanced), or lost (suppressed) during the transition to DE **(Figure 3J-K)**, SI lineage formation **(Figure 3L-M)**, and ileal patterning **(Figure 3N-O)** (log2FC > 2.5, padj < 0.05 by Wald test; DESeq2). We found that a much greater proportion of enhancers, relative to promoters, exhibit significantly altered levels of activity (either gain or loss) across all stage transitions. In contrast, enhancers and promoters were roughly equally represented among those TREs that were unchanged in activity across stage transitions **(Figure 3K, M, O)**. This finding underscores the importance of enhancer dynamics during SI fate acquisition.

### Identification of genes associated with high a density of nearby enhancers that emerge during SI lineage formation or ileal patterning

Next we developed an analysis pipeline to further investigate the enhancers and associated genes that emerge during different stages of the directed differentiation **(Figure 4A)**. We focused on ‘Duo-specific enhancers’ and ‘Ile-specific enhancers’, which are the enhancers that gain activity during Duo and Ile spheroid formation, respectively (**Figure 3M, O**). We then determined the density of stage-specific enhancers for every actively transcribed gene (TPM > 50 in the stage of interest) by counting stage-specific enhancers within a 200 kb window centered on the TSS **(Figure 4A)**. The genes that are associated with the stage-specific enhancers and also have significant activation in both ChRO-seq and RNA-seq are eventually defined as stage-specific enhancer linked genes **(Figure 4A)**.

**Figure 4.**
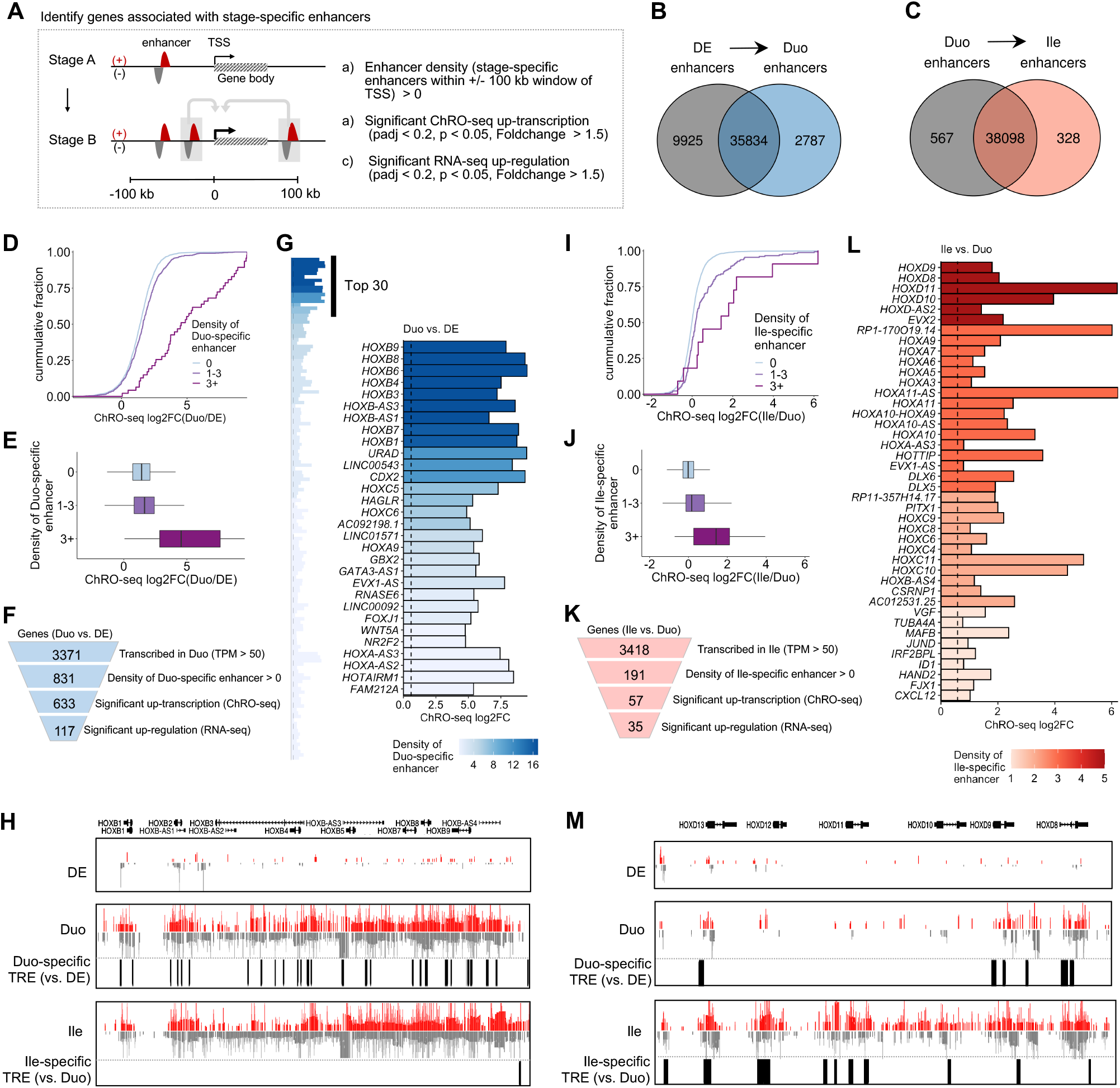
Identification of genes associated with a high density of nearby enhancers that emerge during SI lineage formation or ileal patterning. (A) The strategy for identifying genes associated with stage-specific enhancers. (B) Venn diagram showing stage-specific and shared enhancers between DE and Duo spheroids. Duo-specific enhancers (n = 2787) represent enhancers emerging during SI lineage formation. (C) Venn diagram showing stage-specific and shared enhancers between Duo and Ile spheroids in the event of ileal patterning. Ile-specific enhancers (n = 328) represent enhancers emerging during ileal regional patterning. (D) Cumulative distribution of ChRO-seq fold change in genes grouped into three different categories of Duo-specific enhancer density during the transition to Duo spheroids. (E) Boxplot of ChRO-seq fold change in genes grouped into three different categories of Duo-specific enhancer density during the transition to Duo spheroids. (F) Identification of genes associated with Duo-specific enhancers (n = 117). (G) Bar graph showing genes associated with Duo-specific enhancers (left panel). Top 30 genes based on enhancer density are highlighted (right panel). (H) Normalized ChRO-seq signal around the *HOXB* cluster in the stages of DE, Duo, and Ile spheroids. Duo-specific TREs are marked. (I) Cumulative distribution of ChRO-seq fold change in genes grouped into three different categories of Ile-specific enhancer density during the transition to Ile spheroids. (J) Boxplot of ChRO-seq fold change in genes grouped into three different categories of Ile-specific enhancer density during the transition to Ile spheroids. (L) Identification of genes associated with Ile-specific enhancers (n = 35). (L) Bar graph showing genes associated with Ile-specific enhancers. (M) Normalized ChRO-seq signal around the *HOXD* cluster in the stages of DE, Duo, and Ile spheroids. Ile-specific TREs are marked. ChRO-seq study: DE, n = 4; Duo spheroid (Duo), n = 3; Ile spheroid (Ile), n = 3. RNA-seq study: DE, n = 3; Duo, n = 6; Ile, n = 4.

As defined in **Figure 3**, we identified a total of 2,787 Duo-specific and 328 Ile-specific enhancers **(Figure 4B-C)**. We confirmed that genes associated with a greater number of Duo-specific enhancers also exhibit greater increases in transcription in Duo spheroids relative to DE **(Figure 4D-E)**. Similarly, genes associated with a greater number of Ile-specific enhancers also exhibit greater increases in transcription in Ile relative to Duo spheroids **(Figure 4I-J)**. The results of similar analyses focused on hESC and DE are shown in **Supplementary Figure 4**. These observations support the model in which the level of transcriptional activation during each stage transition is strongly associated with the total number of emerging TREs around TSSs.

Regarding the transition from DE to Duo spheroids, we identified 117 genes that are associated with at least one Duo-specific enhancer and are significantly increased in both transcription (ChRO-seq) and steady-state expression (RNA-seq) **(Figure 4F)**. Among these, as expected, *CDX2* is one of the top genes ranked by the density of Duo-specific enhancers **(Figure 4G)**. Notably, even more highly ranked are the HOXB family members **(Figure 4H)**. Regarding ileal regional patterning, we identified 35 genes that are associated with at least one Ile-specific enhancer and are significantly increased in both transcription and steady state-expression in Ile relevant to Duo spheroids **(Figure 4K)**. Among these, the genes associated with the greatest number of Ile-specific enhancers include members of the *HOXA*/*C*/*D* clusters, *FJX1 CSRNP1*, and *HOTTIP* **(Figure 4L-M)**, several of which were also identified as Ile spheroid-specific marker genes **(Figure 2G)**. These analyses together reveal previously undescribed temporal dynamics of the HOX cluster genes during the events of SI lineage formation followed by ileal regional patterning: first, the HOXB cluster is activated during Duo spheroid formation (likely by nearby Duo-specific enhancers), then subsequently the other three HOX clusters are activated during ileal patterning (likely by nearby Ile-specific enhancers) **(Figure 4H, M)**.

### Identification of genes associated with enhancer ‘hotspots’ that emerge SI lineage formation or ileal patterning

It has been shown in previous studies that dense clusters of highly active enhancers (which we refer to as ‘hotspots’) occur nearby to genes that are especially critical for defining cell identity and status [36-38]. We sought to define for the first time the changing landscape of enhancer hotspots in this human model of SI development. To accomplish this, we adapted a previously described methodology [36, 37], which requires ChIP-seq based datasets, to work with ChRO-seq data instead **(Figure 5A)**. By implementing this analysis pipeline with the Duo- and Ile-specific enhancers (defined in **Figure 4B** and **C** respectively), we were able to identify enhancer hotspots formed during SI lineage formation and ileal patterning, respectively.

**Figure 5.**
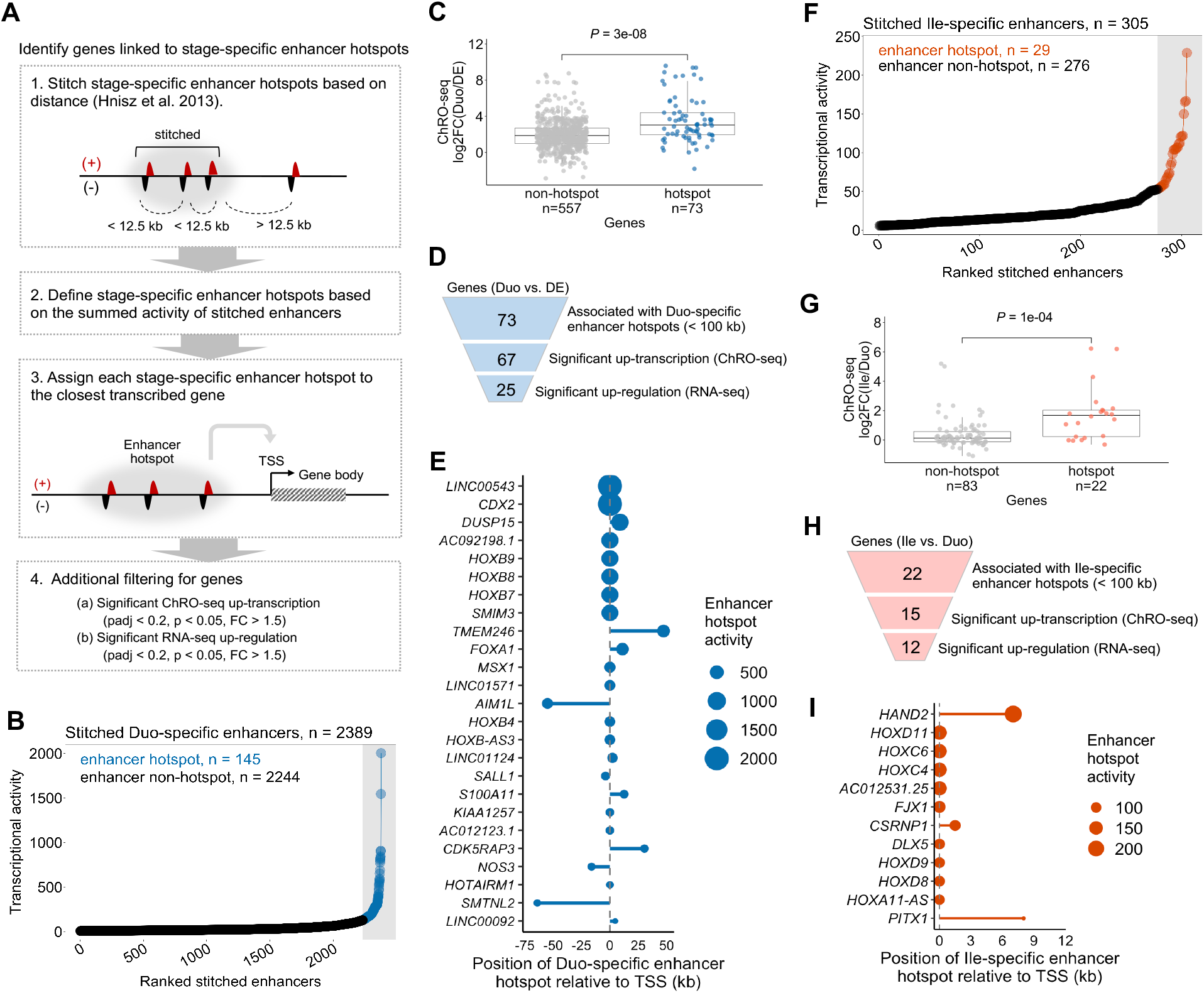
Identification of genes associated with enhancer ‘hotspots’ that emerge during SI lineage formation or ileal patterning. (A) The strategy for identification of stage-specific enhancer hotspots and their associated genes. (B) Duo-specific stitched enhancers are ranked by transcriptional activity (ChRO-seq signal). The stitched enhancers with the highest transcriptional activity are defined as Duo-specific enhancer hotspots (n = 145; blue) and the rest are enhancer non-hotspots (n = 2244; black). (C) ChRO-seq fold change of genes associated with Duo-specific stitched enhancers, non-hotspots vs. hotspots (Wilcoxon rank sum test). (D) Identification of genes associated with Duo-specific enhancer hotspots. (E) Genes associated with Duo-specific enhancer hotspots (n=25). Relative position between enhancer hotspots and transcription start sites (TSSs) of the associated genes are shown. Dot size denotes transcriptional activity of a given DE-specific enhancer hotspot. (F) Ile-specific stitched enhancers are ranked by transcriptional activity (ChRO-seq signal). The stitched enhancers with the highest transcriptional activity are defined as Ile-specific enhancer hotspots (n = 29; red) and the rest are enhancer non-hotspots (n = 276; black). (G) ChRO-seq fold change of genes associated with Ile-specific stitched enhancers, non-hotspots vs. hotspots (Wilcoxon rank sum test). (H) Identification of genes associated with Ile-specific enhancer hotspots. (I) Genes associated with Ile-specific enhancer hotspots (n=12). Relative position between enhancer hotspots and TSSs of the associated genes are shown. Dot size denotes transcriptional activity of a given Duo-specific enhancer hotspot. ChRO-seq study: DE, n = 4; Duo spheroid (Duo), n = 3; Ile spheroid (Ile), n = 3. RNA-seq study: DE, n = 3; Duo, n = 6; Ile, n = 4.

To identify enhancer hotspots and the associated candidate genes that may be critical for SI lineage formation, we first clustered Duo-specific enhancers into ‘stitched enhancers’ **(Supplementary Data 3)**. Among the 2389 stitched enhancers, we identified 145 that exhibit strong enough transcriptional activity to be designated as Duo-specific ‘enhancer hotspots’ **(Figure 5B)**. We assigned each of the Duo-specific stitched enhancers (including non-hotspots and hotspots) to its nearest active gene (TPM > 50 in ChRO-seq) within a 100 kb window on either end of the boundaries of the stitched enhancer. We found that the overall increase in gene body transcription is significantly greater for the set of genes associated with Duo-specific enhancer hotspots compared to those associated with stitched enhancers that are not hotspots **(Figure 5C)**. This finding is consistent with the notion that hotspots exert particularly strong effects on transcription. Among the 73 genes nearest to Duo-specific enhancer hotspots, 25 exhibit highly significant increases in both transcription (ChRO-seq) and steady state expression (RNA-seq) in Duo spheroids relative to DE **(Figure 5D-E)**. Many of these were defined earlier as SI or Duo-specific markers (e.g., *CDX2, FOXA1, HOXB* cluster, *SALL1* and *LINC00543*) **(Figure 2)**.

We next performed a similar analysis to identify enhancer hotspots and nearby genes associated with ileal regional specification. Among 305 stitched enhancers formed by Ile-specific enhancers (relative to Duo spheroids) **(Supplementary Data 3)**, we defined 29 Ile-specific enhancer hotspots **(Figure 5F)**. We assigned each of the Ile-specific stitched enhancers (including non-hotspots and hotspots) to its nearest active gene (TPM > 50 in ChRO-seq) within a 100 kb window on either end of the boundaries of the stitched enhancer. Similar to the observation made for the transition from DE to Duo spheroids, the genes nearest to Ile-specific enhancer hotspots exhibit a significantly greater increase in transcription compared to those associated with non-hotspot stitched enhancers **(Figure 5G)**. Among the 22 genes nearest to Ile-specific enhancer hotspots, 12 exhibit highly significant increases in both transcription (ChRO-seq) and steady-state expression (RNA-seq) in Ile relative to Duo spheroids **(Figure 5H)**. These genes include *HAND2, HOXC/D* family members, *FJX1, DLX5, CSRNP1* and *PITX1* **(Figure 5I)**, many of which were also identified earlier as Ile-specific markers **(Figure 2)**. Similar analyses were performed for the hESC and DE stages and the results are summarized in **Supplementary Figure 5**.

### Identification of candidate TF drivers and their networks relevant to SI lineage formation or ileal patterning

To discover putative key TF drivers and their cistromes (genome-wide set of TF targets) in this model, we first used the tool HOMER to determine motifs enriched in each set of stage-specific enhancers and then identified genes associated enhancers that harbor specific motifs of interest **(Figure 6A)**. Expectedly, HOMER analyses of hESC- and DE-specific enhancers revealed binding sites of POU5F1 and SMAD2 to be the most enriched motifs, respectively **(Supplementary Figure 6A-D)**. We next performed motif enrichment analyses on Duo- **(Figure 6B)** and Ile-specific enhancers **(Figure 6D)**.

**Figure 6.**
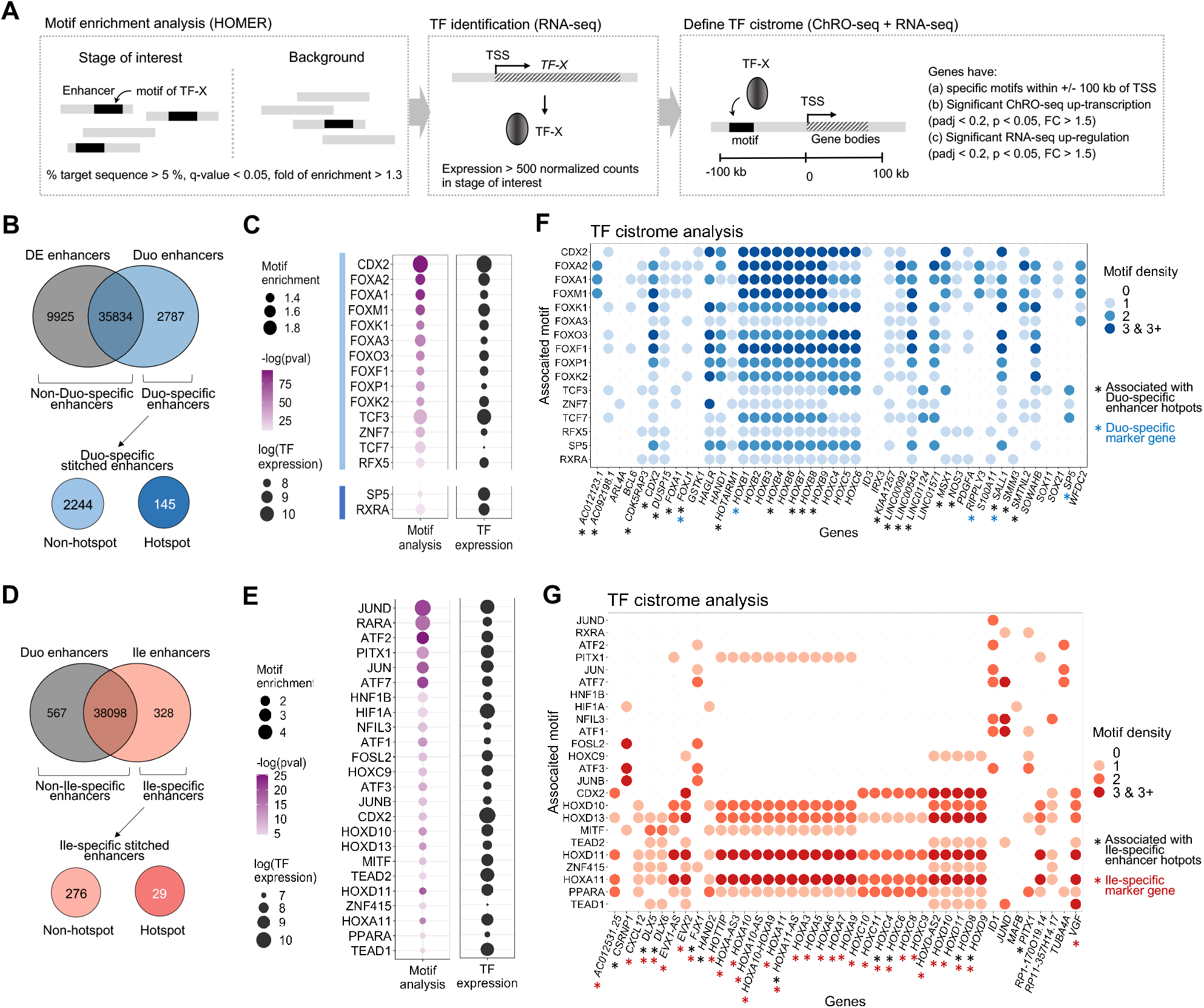
Identification of candidate TF drivers and their networks relevant to SI lineage formation or ileal patterning. (A) The strategy for identifying candidate TFs and their cistromes associated with each stage transition of the model. (B) TF motif enrichment analyses were performed in Duo-specific enhancers (n = 2787) relative to non-Duo-specific enhancers (n = 45,759) as well as in Duo-specific enhancer hotspots (n = 145) relative to non-hotspots (n = 2,244). (C) Motifs significantly enriched in Duo-specific enhancers (highlighted by light blue bar) or in Duo-specific enhancer hotspots (highlighted by dark blue bar). The steady-state expression (RNA-seq) of the corresponding TFs are also shown. (D) TF motif enrichment analyses were performed in Ile-specific enhancers (n = 328) relative to non-Ile-specific enhancers (n = 38,674) as well as in Ile-specific enhancer hotspots (n = 29) relative to non-hotspots (n = 276). (E) Motifs significantly enriched in Ile-specific enhancers. The steady-state expression (RNA-seq) of the corresponding TFs are also shown. (F) Bubble plot showing genes associated with active binding motifs of TFs that exhibit an overall enrichment of binding motifs in Duo-specific enhancers (defined in C). See also **Supplementary Figure 6E** for full gene list. (G) Bubble plot showing genes associated with active binding motifs of TFs that exhibit an overall enrichment of binding motifs in Ile-specific enhancers (defined in E). ChRO-seq study: DE, n = 4; Duo spheroid (Duo), n = 3; Ile spheroid (Ile), n = 3. RNA-seq study: DE, n = 3; Duo, n = 6; Ile, n = 4.

As expected, among the 2,787 Duo-specific enhancers relative to non-Duo specific enhancers **(Figure 6B)**, we detected significant enrichment for the binding motifs of CDX2 and many forkhead box TFs **(Figure 6C)**. We also performed the same analysis with Duo-specific enhancer hotspots relative to non-hotspots **(Figure 6B)** and observed significant enrichment for binding motifs of SP5 and RXR **(Figure 6C)**, the former of which is defined as a Duo-specific gene marker **(Figure 2G)**, but neither of which have been associated previously with early SI development. Genes that are associated with Duo-specific enhancers harboring motifs of one or more of these TFs and that are significantly increased in transcription (ChRO-seq) and steady-state expression (RNA-seq) are shown in **Figure 6F (see full gene list in Supplementary Figure 6E)**. The TF cistrome analysis suggests that CDX2 is a prominent transcriptional activator of the *HOXB* cluster **(Figure 6F)**, which is consistent with previous observations [8]. Moreover, this analysis reveals candidate TF activators of the gene *CDX2*. While *CDX2* locus has a CDX2 motif in its promoter region in Duo-spheroids (data not shown) and is thus likely subject to autoregulation as previously described [39], Duo-specific enhancers that are associated with the *CDX2* locus do not harbor any binding motifs for CDX2 and instead have binding motifs for other TFs such as FOXA1 and FOXO3 **(Figure 6F)**. More importantly, through this analysis we were able to identify candidate TF drivers of SI/Duo spheroid marker genes that do not appear to be a part of the CDX2 cistrome. For example, *WFDC2* and *SALL1* (defined earlier as a marker gene of SI and Duo spheroids respectively) are associated with FOXA1 and FOXO3 motifs; *SP5* (defined earlier as a marker gene of Duo spheroid) is associated with the TCF3 motif **(Figure 6D)**.

We next performed a similar analysis for ileal regional patterning. Among the 328 Ile-specific enhancers relative to non-Ile specific enhancers **(Figure 6D)**, we detected significant enrichment for the binding motifs of several key TF families including CDX2, HOXC9, HOXA11 and PITX1 **(Figure 6E)**, the last of which is itself associated with Ile-specific enhancer hotspots **(Figure 5I)**. We also performed the same analysis with Ile-specific enhancer hotspots relative to non-hotspots but did not observed significant enrichment for binding motifs (data not shown). Genes that are associated with Ile-specific enhancers harboring motifs of one or more of these TFs and that are significantly increased in transcription (ChRO-seq) and steady-state expression (RNA-seq) are shown in **Figure 6G**. This analysis led to several new observations. Firstly, CDX2 likely plays a role in both Duo spheroid formation and ileal regional patterning, but apparently through distinct cistromes **(Figure 6F and G)**. Notably, the activation of *HOXD* and *HOXC* (especially *HOXC8* and *9*) loci is associated with Ile-specific TREs harboring the CDX2 motif **(Figure 6E)**. While the HOXB cluster is induced in Duo spheroids, it remains highly transcribed in Ile spheroids, likely due to the fact that the nearby TREs with CDX2 motifs that are activated during Duo spheroid formation are retained during ileal regional patterning **(Figure 4H, M)**. Secondly, we found that the activation of HOXA, C, and D families are associated with overlapping but distinct TF regulators. Specifically, while HOXA11 seems to act on all four *HOX* clusters, PITX1 is associated only with the *HOXA* cluster and HOXC9 is preferentially associated with the *HOXD* cluster **(Figure 6G)**. Finally, this analysis also reveals other TF cistrome networks that do not target *HOX* genes during ileal patterning **(Figure 6G)**. For example, *CSRNP1, FJX1* and *ID1* loci are associated with motifs of AP-1 (e.g. JUNB and FOSL2) and/or ATF factors. **(Figure 6G)**.

### The HOX cluster dynamics observed in the directed differentiation model of SI is present in the early developmental stages of primary human fetal SI tissue

Across various analyses in this study we repeatedly observed a previously undescribed dynamics of HOX gene activation across SI lineage formation to ileal patterning. Notably, ChRO-seq data reveals strong transcriptional activation of the *HOXB* cluster (particularly HOXB1) upon acquisition of SI identity (Duo spheroids) and the sequential transcriptional activation of members of *HOXA, C, D* clusters during ileal regional patterning (Ile spheroids) **(Figure 7A)**, which as we showed previously is also reflected in the pattern of emergence of nearby enhancers and enhancer hotspots **(Figure 4,5)**. This unique HOX cluster dynamics is largely reflected at the level of steady-state expression (RNA-seq) also **(Figure 7B)**, except for the 3’-most *HOXA* genes (e.g. *HOXA1, A2*) and 5’-end HOXB genes (e.g. *HOXB7, B9*) **(Figure 7A-B)**, which may be subject to post-transcriptional regulation.

**Figure 7.**
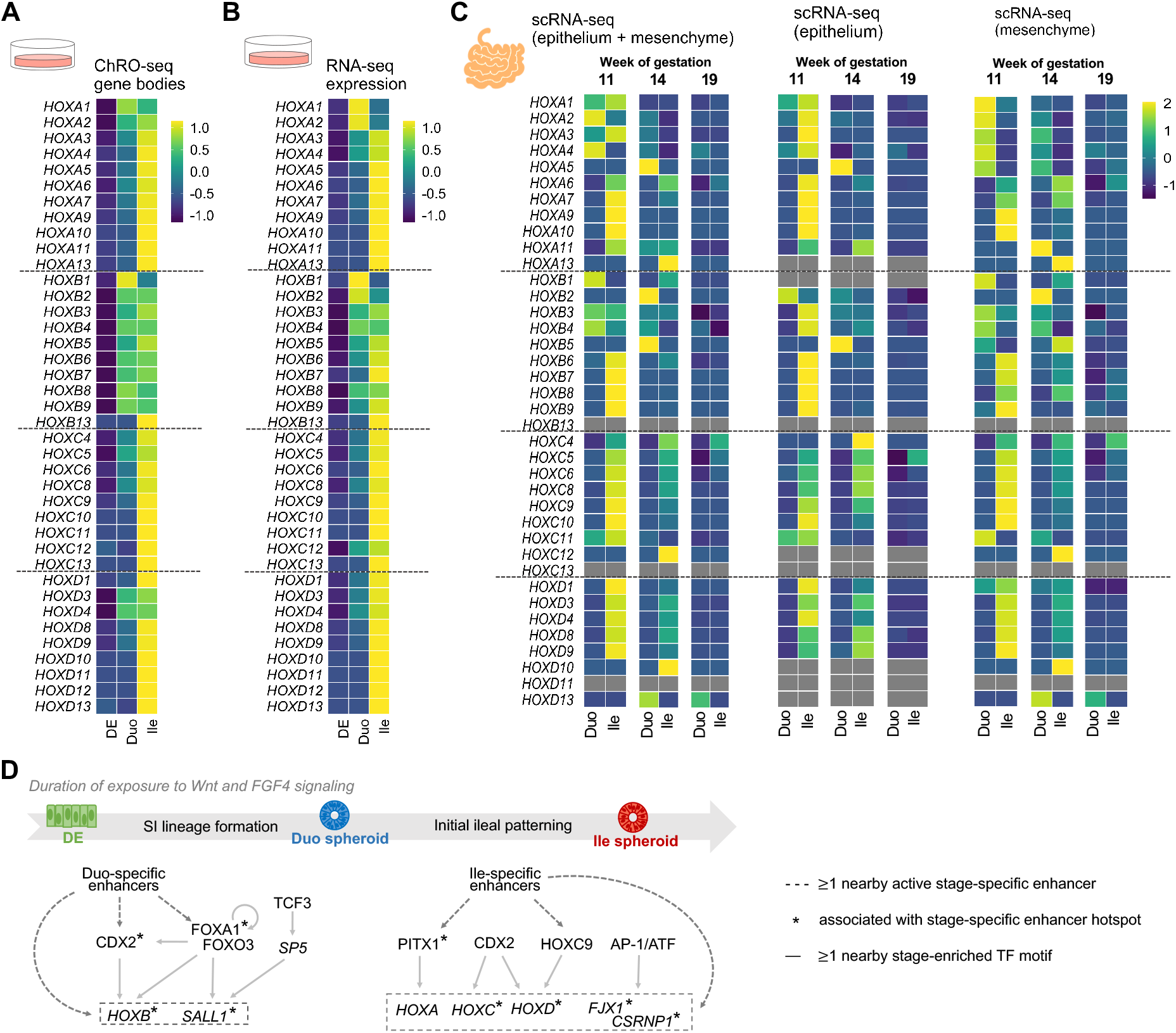
The HOX cluster dynamics observed in the directed differentiation model of SI is present in the early developmental stages of primary human fetal SI tissue. (A) Heatmap showing the changing transcriptional pattern of HOX genes (ChRO-seq) across the stages of DE, Duo spheroids, and Ile spheroids *in vitro*. (B) Heatmap showing changing steady-state expression pattern of HOX genes (RNA-seq) across stages of DE, Duo spheroids, and Ile spheroid *in vitro*. (C) Heatmap showing changing expression pattern of HOX genes (scRNA-seq) in primary human fetal Duo and Ile tissues at week 11, 14 and 19 of gestation. The expressional level of each HOX gene is the average from the fraction that has both epithelial and mesenchymal cell types (left), mesenchymal cell only (middle), or epithelial cell only (right). (D) Proposed model of transcriptional programming in SI lineage formation and ileal regional patterning during human SI development. The genes activated in the relevant stages and the types of regulatory relationships identified through ChRO-seq analyses are indicated. ChRO-seq study: DE, n = 4; Duo spheroid (Duo), n = 3; Ile spheroid (Ile), n = 3. RNA-seq study: DE, n = 3; Duo, n = 6; Ile, n = 4. scRNA-seq data: matched Duo and Ile tissues from 1 human fetal subject per gestational time point.

To determine whether the differences in *HOX* gene expression between early Duo and Ile extends beyond this *in vitro* directed differentiation model, we mined the single cell RNA-seq (scRNA-seq) dataset of primary human Duo and Ile tissue at very early stages of fetal development [13, 40, 41]. Specifically, we focused on matched Duo and Ile fetal tissue at week 11, 14 and 19 of gestation. It is important to note that these developmental time points, though considered as early/mid gestation stage, represent time points much later than what is reflected in our much earlier stage SI spheroids, and therefore exhibited weaker overall expression of *HOX* genes. Since the SI spheroids generated from the directed differentiation method are mainly comprised of epithelial cells but also contain some mesenchymal cells, we analyzed both cell types from the scRNA-seq data, together and separately, in order to examine the *HOX* dynamics. At week 11 of gestation, we found that the *HOXB* cluster is active in both human fetal Duo and Ile cells (epithelium plus mesenchyme), with *HOXB1* particularly enriched in Duo cells and 5’ *HOX* genes of all four clusters enriched in Ile cells **(Figure 7C)**. This distinct HOX pattern between primary human fetal Duo and Ile tissues is consistent with our observations in the Duo and Ile spheroids generated from the directed differentiation method. Interestingly, this signal is present only at week 11 and dissipates as the developmental process proceeds (week 14 and 19) **(Figure 7C)**. The scRNA-seq analysis also offers the new insight that the enrichment of *HOXB1* in primary fetal Duo tissue is mainly driven by the mesenchymal fraction, whereas the activation of 5’ HOX genes of all four clusters in Ile tissue occurs in both epithelial and mesenchymal cells **(Figure 7C)**. Overall, this analysis suggests that the Duo and Ile spheroids generated from the directed differentiation model appear to capture the region-specific HOX features that are transiently present in the developing human SI.

Beyond the finding of the previously undescribed temporal HOX cluster patterns highlighted here, the ChRO-seq analyses in this study provide a comprehensive characterization of chromatin regulatory dynamics, enhancer activity profiles, and transcriptional programs that are associated with human SI lineage formation and regional patterning. A working model is provided in **Figure 7E**.

## Discussion

By leveraging the recently developed ChRO-seq technology, we generated for the first time ever comprehensive chromatin regulatory landscapes across the different stages of directed differentiation from hPSC to SI spheroids with regional specification. Specifically, we defined: (1) early marker genes that label SI lineage formation and regional specification, (2) the map of <u>active</u> regulatory elements (promoters and enhancers) as well as enhancer hotspots associated with SI lineage formation and ileal patterning, and (3) candidate key TF drivers and their cistromes relevant to these critical developmental events. These findings motivate future experiments to investigate the functional role of candidate regulators and assess the sufficiency and necessity of the associated enhancers and enhancer hotspots for controlling transcriptional networks relevant to early human SI development and patterning.

We identified for the first time *HES4* as marker gene of SI spheroids irrespective of regional identity in the context of gut development; one likely reason for this is that it is absent in the genome of the mouse [42], the model organism most used previously to study gut development. Moreover, the marker genes that we found to specifically label Duo identity include *SP5* and *SALL1*, the latter of which has been linked to Townes-Brocks syndrome with developmental malformation of multiple organs including the limb, kidney and gastrointestinal tract [43, 44]. While *SP5, SALL1* and other marker genes of Duo identity were reported to enhance Wnt signaling [45, 46], we identified a distinct set of marker genes that specifically label Ile identity (e.g. *HAND2, FJX1* and *CSRNP1*) that are also known to be downstream of Wnt signaling. *Fjx1* has been shown to function through the Wnt/planar cell polarity (PCP) pathway to determine anterior-posterior axis in *Drosophila* [47, 48] and is expressed in the epithelium of murine developing gut [49]. *Csrnp1* is expressed in neural crest progenitors in a Wnt1/β-catenin-dependent manner and acts as a critical effector of Wnt signaling for driving neural crest formation [50]. Together, these observations indicate that different sets of Wnt signaling effector genes are likely induced upon different durations of exposure to the Wnt-activating supplements during directed differentiation. Future investigation of the functional roles of these genes in the context of early gut development is warranted. The present study also uncovers the associated enhancers (or enhancer hotspots) and TF motifs of these marker genes, which may elucidate the potential mechanisms through which these genes are regulated during the establishment of SI identity and regional specification.

Although HOX genes have been reported to play important roles in proximal-distal axis patterning in the gut, our study uncovers previously undescribed dynamics of HOX cluster genes in the directed differentiation model of human SI development, unveiling the complexity of HOX biology in the control of small intestinal regionalization. The early literature has observed that 3’ HOX genes are relevant to proximal gut and 5’ HOX genes are restricted to distal gut (known as ‘spatial collinearity’ of HOX genes) [51, 52]; however, our study shows that HOX genes do not simply follow this collinear pattern. We found that the activation of the HOXB cluster first occurs during the formation of Duo spheroids and the other HOX clusters (HOXA, C and D) are activated subsequently during the formation of Ile spheroids. Through mining of a novel scRNA-seq dataset [13, 40, 41], we also showed that this distinct HOX behavior between Duo and Ile regions is present in primary human fetal SI tissue. Through ChRO-seq analyses, we also provided additional insights into HOX biology by elucidating candidate TF drivers and associated enhancers and enhancer hotspots, which merits detailed investigation.

Our ChRO-seq analysis mapped the key transcriptional networks associated with SI lineage formation or regional patterning. For example, we found that binding sites of FOXA1 and FOXO3 are enriched in active enhancers that emerge only during SI identity acquisition, including those nearby *CDX2*, which encodes the well-known master TF regulator of SI development. Notably, the *FOXA1* gene locus is actively up-transcribed during the transition from definitive endoderm to SI spheroids and is associated with an enhancer hotspot that emerges during this process. FOXA1 has been reported to regulate allocation of murine SI secretory lineage [53] and proper gene expression in murine intestinal epithelium [54]. Recently, dysregulation of intestinal *FOXA1* also has been linked to necrotizing enterocolitis in infants [55]. The observations made in this study as well as previous reports together underscores the functional importance of FOXA1 in human gut development. Also, our TF cistrome analyses indicate that CDX2 is involved in not only in initial SI lineage establishment but also the subsequent event of ileal specification, but evidently through distinct cistromes. One specific example is the association between the *HOXC9* locus and the nearby Ile-specific active enhancers containing CDX2 binding motifs, which is consistent with a previous study showing a remarkable downregulation of *Hoxc9* in distal but not in proximal intestine of *Cdx2* null mice [2]. Recent evidence also demonstrates that CDX2 regulates developing and fully mature gut through targeting distinct gene sets in mice [8] and that CDX2 is involved in patterning both intestinal epithelial and mesenchymal cells in the hPSC-based directed differentiation model [13]. Our ChRO-seq findings pertaining to CDX2 may further elucidate how CDX2 functions through a stage- or cell type-dependent manner during early human gut development. Beyond CDX2, other candidate TFs associated with the establishment of ileal identity are worthy of future investigation. One notable example is PITX1, the ortholog of which has been reported previously in mouse models to be expressed in the distal SI region and to be involved in cecum budding during murine development [56]. However, PITX1 has not been previously appreciated in the context of ileal regional specification.

The use of the state-of-art *in vitro* model of human SI development is a critical feature of this study given the lack of other existing platforms or resources that are readily available for the study of very early developmental time points pertaining to SI lineage formation and regional patterning. It is also important given the growing appreciation for cross-species differences in developmental processes at the molecular level [57-60]. Furthermore, the genome-wide characterization of the chromatin regulatory landscape by ChRO-seq in this model system generates valuable translational knowledge for a better understanding of the transcriptional programming underlying early SI development in humans. Our analysis pipeline can also be applied to other hPSC-derived organoid systems to gain insights into dynamic transcriptional programming and chromatin status during early stages of human organogenesis. Most importantly, the identification of the candidate drivers in the present study may ultimately improve methods of generating therapeutic replacement SI as well as molecular therapies for children and possibly adult patients.

## Methods

### Directed differentiation of hESCs

Differentiation of H9 hESCs and organoids was performed as previously published, with minor modifications [27, 61]. Briefly, the endoderm was generated by treatment of activin A (100 ng/ml) for 3 consecutive days in Roswell Park Memorial Institute 1640 (RPMI-1640) media supplemented with 0% (v/v), 0.2% (v/v) and 2.0% (v/v) Hyclone defined fetal bovine serum (dFBS). The endoderm cultures then received daily treatments of FGF4 (500 ng/mL) and CHIR99021 (2 μM) for next 10 days. The intestinal spheroids representing fetal duodenum and ileum were collected on day 5 and day 10, respectively [28]. Subsets of spheroids collected at day 5 and day 10 were then cultured in Matrigel with the previously defined intestine growth media[28] for 28 days in order to mature into organoids. The resulting organoids were then prepared for cell sorting to purify EPCAM+ cell population (epithelial fraction) and the sorted cells were processed for RNA-seq library preparation. The cells generated at the stages of hESC, endoderm, duodenal spheroids and ileal spheroids were subject for ChRO-seq and RNA-seq library preparation.

### Fluorescence activated cell sorting (FACS)

Methods for organoid dissociation into single cells followed by selection of the epithelial component with FACS, were based on previously described procedures[62]. All solutions, including overnight pretreatment of the organoid cultures, contained 10μM Y27632 (Tocris). Matrigel was digested for 30 minutes with cold 4mM EDTA-DPBS and organoids were washed 4X with cold DPBS. Structures were enzymatically dissociated into single cells using the Tumor Dissociation Kit (human) (Miltenyi Biotec) with a gentleMACS™ Octo Dissociator (with heaters; Miltenyi Biotec) for 50 minutes at 37°C. The cell suspension was then washed with 0.5% BSA-2mM EDTA-DPBS over a succession of cell strainers, 100μm, 40μm (Corning) and 20μm (CellTrics) and centrifuged for 5 minutes at 500xg. Cells were labeled with EpCAM phycoerythrin (PE)-conjugated antibody (BioLegend) and an EpCAM isotype-PE control (BioLegend), and were sorted in 0.1% BSA 2mM EDTA-DPBS on a MoFlo Astrios 1 (Beckman Coulter; Brea, California) instrument at the University of Michigan BRCF Flow Cytometry Core facility. Events were first selected with light-scatter and doublet discrimination gating, followed by exclusion of dead cells using 1μM DAPI dilactate (Molecular Probes). EpCAM-PE(+)/DAPI(-) cells were sorted into cold Advanced DMEM/F12 (Invitrogen). Collected cells were reanalyzed for a purity-check and showed greater than 89% viable and 98% EpCAM-PE(+) events. Cells were pelleted at 500xg for 5 minutes and flash-frozen for subsequent RNA isolation.

### Procr-mGFP-2A-LacZ mouse line and staining

The Procr-mGFP-2A-LacZ mouse line was generated by knocking in a cassette of mGFP-2A-LacZ behind start codon of the *Procr* gene [32]. To perform X-gal staining, embryos were isolated and washed in cold PBS followed by incubation in ice-cold fixative (30 mins up to E10.5, or 50 mins for E11.5-E13.5) on a rocking platform. Fixative contains 37% formaldehyde, 25% glutaraldehyde, 10% NP-40 all dissolved in PBS. The fixative was removed and the whole embryo washed twice in PBS for 20 mins at room temperature on a rocking platform. The β-galactosidase substrate (5 mM K3FE(CN)6, 5 mM K4Fe(CN)6·3H2O, 2 mM MgCl2, 0.02% NP40, 0.1% sodium deoxycholate and 1 mg ml–1 X-gal in PBS) was then added and the tissues incubated in the dark overnight at room temperature. The substrate was removed and the tissues washed twice in PBS for 20 min at room temperature on a rocking platform. Embryos were serially dehydrated using glycerol, and was stored in 80% glycerol at 4°C. Whole-embryo images were captured using Olympus SZX16.

### Chromatin isolation

The chromatin isolation for ChRO-seq library preparation was performed as previously described [20, 63]. Briefly, chromatin was isolated from the cells with 1× NUN buffer [0.3 M NaCl, 1 M Urea, 1% NP-40 (w/v), 20 mM HEPES, pH 7.5, 7.5 mM MgCl2, 0.2 mM EDTA, 1 mM DTT, 20 units per mL SUPERase In RNase Inhibitor (Life Technologies, AM2694), 1× Protease Inhibitor Cocktail (Roche, 11 873 580 001)] and incubation at 12 °C on a ThermoMixer for 30 min. Samples were centrifuged at 12,500xg for 30 min at 4 °C. The chromatin pellet was washed 3 times with 1 mL 50 mM Tris-HCl, pH 7.5, supplemented with 40 units per ml RNase inhibitor. Chromatin storage buffer (50 mM Tris-HCl, pH 8.0, 25% glycerol (v/v), 5 mM Mg(CH3COO)2, 0.1 mM EDTA, 5 mM DTT, 40 units per ml RNase inhibitor) was added to each sample. The samples were loaded into a Bioruptor and sonicated to get the chromatin into suspension. Samples were stored at −80 °C before proceeding to ChRO-seq library preparation.

### ChRO-seq library and sequencing

After chromatin isolation, the ChRO-seq library preparation closely followed the protocol described previously [20, 64]. Briefly, chromatin from at least 1 × 10^6^ cells per sample in chromatin storage buffer was mixed with an equal volume of 2×chromatin run-on buffer [10 mM Tris-HCl, pH 8.0, 5 mM MgCl2,1 mM DTT, 300 mM KCl, 400 μM ATP (NEB, N0450S), 40 μM Biotin-11-CTP (Perkin Elmer, NEL542001EA), 400 μM GTP (NEB, N0450S), 40 μM Biotin-11-UTP (Perkin Elmer, NEL543001EA), 0.8 units per μL RNase inhibitor, 1% (w/v) Sarkosyl (Fisher Scientific, AC612075000)]. The run-on reaction was incubated at 37 °C for 5 min. The reaction was stopped by adding Trizol LS (Life Technologies, 10296-010) and pelleted with GlycoBlue (Ambion, AM9515) to visualize the RNA pellet. The RNA pellet was resuspended in diethylpyrocarbonate (DEPC)-treated water and heat denatured at 65 °C for 40 s. In the present study, base hydrolysis of RNA was performed by incubating RNA with 0.2N NaOH on ice for 4 min. Nascent RNA was purified by binding streptavidin beads (NEB, S1421S) before and in between the following procedures: (1) 3′ adapter ligation with T4 RNA Ligase 1 (NEB, M0204L), (2) 5′ de-capping with RNA 5′ pyrophosphohydrolase (RppH, NEB, M0356S), (3) 5′ end phosphorylation using T4 polynucleotide kinase (NEB, M0201L), (4) 5′ adapter ligation with T4 RNA Ligase 1 (NEB, M0204L). The resulting RNA fragments were used for a reverse transcription reaction using Superscript III Reverse Transcriptase (Life Technologies, 18080-044) to generate cDNA. cDNA was then amplified using Q5 High-Fidelity DNA Polymerase (NEB, M0491L) to generate the ChRO-seq libraries. Libraries were sequenced (5’ single end; single-end 75x) using the NextSeq500 high-throughput sequencing system (Illumina) at the Cornell University Biotechnology Resource Center. **Supplementary Data 1** provides the mapping statistics of the ChRO-seq experiments.

### Total RNA isolation, mRNA-seq library and sequencing

Total RNA was isolated using the Total Purification kit (Norgen Biotek, Thorold, ON, Canada). High Capacity RNA to cDNA kit (Life Technologies, Grand Island, NY) was used for reverse transcription of RNA. Libraries were generated using the NEBNext Ultra II Directional Library Prep Kit (New England Biolabs, Ipswich, MA) and subjected to sequencing (single-end 92x) on the NextSeq500 platform (Illumina) at the Cornell University Biotechnology Resource Center. At least 80M reads per sample were acquired.

### Mapping sequencing reads

In the ChRO-seq studies, the publicly available pipeline [33] was used to align ChRO-seq reads. Since the libraries were prepared using adapters that contained a molecule-specific unique identifier (first 6 bp sequenced), the PCR duplicates were first removed using PRINSEQ lite. Adapters were trimmed from the 3′ end of remaining reads using cutadapt with a 10% error rate. Reads were mapped with the Burrows-Wheeler Aligner (BWA) to the human reference genome hg38 plus a single copy of the Pol I ribosomal RNA transcription unit (GenBank U13369.1). The location of active RNA polymerase was represented by a single base that denotes the 3′ end of the nascent RNA, which corresponds to the position on the 5′ end of each sequenced read. Mapped reads were converted to bigwig format using BedTools and the bedGraphToBigWig program in the Kent Source software package. For visualization purpose, bigwig files from identical stages were merged and normalized to a total signal of 1×10^6^. In the RNA-seq studies, reads were mapped to human genome hg38 using STAR (v2.5.3a) [65] and transcript quantification was performed using Salmon (v0.6.0) [66] with GENCODE release 25 transcript annotations. The expression (RNA-seq) levels of genes were normalized using DESeq2 [67]. All the samples except an Ile HIO sample had < 80% mapping rates. Although the Ile HIO sample had an unfavorable mapping rate, it was able to show elevated levels of Ile-associated regional markers compared to the Duo HIO sample **(Supplementary Figure 3A)**.

### ChRO-seq quantification of gene loci and promoter transcription activity

Gene definitions were obtained from GENCODE v25 annotations. To quantify transcription activity of gene loci, ChRO-seq signals present in annotated gene bodies were used with exclusion of reads within 500 base downstream of transcription start site (TSS) to avoid bias generated by the RNA polymerase pausing at the promoters. Genes with gen body < 1000 base were excluded from all the gene body related analysis, given that genes with short gene bodies are likely biased when excluding the pause peak. The ChRO-seq reads were normalized by the length of gene bodies to transcripts per million (TPM). To determine promoter activity of genes, stranded ChRO-seq signals at the promoter proximal region (500 bps upstream and 200 bps downstream of TSS) were used. The promoter regions that have significant changing patterns were defined using the following criteria: average TPM > 5 and padj < 0.05 across all the stages by likelihood ratio test (DESeq2). The promoter regions that have significant changing patterns and are annotated as ‘protein-coding’ gene type were further subject to clustering analysis (‘degPatterns’ in R platform).

### Differential expression and pathway analyses of genes

The differential analysis of gene bodies in ChRO-seq data was performed using DESeq2 package [67]. For all the analyses except stage-specific marker analysis, the normalized levels of transcription or expression, foldchange and the statistic filtering were based on the DESeq2 analysis including only the two stages in a comparison. The pathway enrichment analyses with subsets of genes were assessed using Enrichr [68].

### Identification of stage-specific marker genes

Firstly, genes that have ChRO-seq signal in gene bodies (excluding the first 500 bps downstream of the TSS) greater than 50 TPM in the stage of interest are identified and filtered according to padj < 0.2, p < 0.05, fold change > 1.5 compared to all the other stages (Wald test; DESeq2). Secondly, genes that have RNA-seq signal greater than 100 base mean units are identified and filtered according to padj < 0.2, p < 0.05, fold change > 1.5 compared to all the other stages (Wald test; DESeq2). Thirdly, the output from these two analyses are intersected to arrive at final gene lists for stage-specific markers. To identify genes that label SI spheroids irrespective of regional identity, analyses were done using the same criteria except the fold change criterion. The genes that have short gene bodies (< 1000 bp) or annotated under the pseudogene category are excluded from this analysis.

### TRE identification, annotation, categorization and comparison with other datasets

The active transcriptional regulatory elements (TREs) were identified by dREG tool [33]. The annotation of the identified TREs was defined using *annotatePeaks*.*pl* function (genome = hg38) of HOMER package [69] based on GENCODE v25 annotations. To compare the TRE landscape of SI spheroids with developing and mature SI in humans, DNase-seq datasets from the Roadmap Epigenomics Project were used. Specifically, first, the files E085-DNase.hotspot.fdr0.01.peaks.v2.bed (fetal SI) and E109-DNase.hotspot.fdr0.01.peaks.v2.bed (adult SI) were downloaded in hg19, converted to hg38 using the USCS liftOver tool, and used to define DNase-based open chromatin regions unique to fetal and adult SI using the bedtools intersect function (at least 1 base overlapping). The fetal and adult SI-specific DNase open chromatin regions were further intersected with TREs present in SI spheroids using bedtools intersect function (at least 1 base overlapping). To assign active TREs as promoters, the TREs with at least 1 base overlapping with the window of -1000 base and +200 base of annotated TSSs were defined as proximal TREs, or promoters. The rest of the active TREs were defined as distal TREs, or enhancers.

### Stage-specific TRE density analysis

The stage-specific TREs were defined as TREs of which the ChRO-seq intensity is significantly higher in a stage of interest relative to a comparative stage (padj < 0.05 and log_2_fold change > 2.5 by DESeq2) [67]. The ChRO-seq intensity was determined by the sum of the un-normalized ChRO-seq reads from both strands within a TRE region. The density of stage-specific TREs was determined by the number of TREs present within the window of +100 kb and -100 kb around the TSS for all the genes which are actively transcribed in the stage of interest (TPM > 50 in the ChRO-seq study). The genes which are associated with stage-specific TREs are defined using the following criteria: (1) genes have density of stage-specific TRE > 0, (2) genes are actively transcribed (TPM > 50 in the ChRO-seq) and expressed (base mean > 100 in the RNA-seq) in the stage of interest, and (3) the genes are significantly uptranscribed and upregulated (padj < 0.2, p < 0.05, fold change > 1.5 in both ChRO-seq and RNA-seq by DESeq2) in the stage of interest relative to a comparison stage.

### Identification of de novo stage-specific enhancer hotspots and the associated genes

The stage-specific active distal TREs (enhancers) were used in the enhancer hotspot analysis. The enhancer hotspots in this study were identified by the criteria similar (with slight modifications) to the studies describing ‘super-enhancers’[36, 37]. Briefly, the stage-specific enhancers in proximity of distance (< 12.5kb) were stitched. For each of the stage-specific stitched enhancers, the transcription activity was determined by the sum of un-normalized ChRO-seq signals from both strands of each individual enhancer. To further identify stage-specific enhancer hotspots, a tangent line was applied the stage-specific stitched enhancers and they were ranked based on their transcriptional activities in a plot. The ones above the tangent line in the analysis were defined as stage-specific enhancer hotpots. To identify the genes which are associated with stage-specific stitched enhancers, a given stage-specific stitched enhancer is assigned to the gene of which the transcription in the gene body is active (TPM > 50 in the matching stage) and the TSS is closest to the border of the enhancer hotspot region.

### Transcription factor binding motif enrichment analysis

HOMER tool [69] was used to determine enrichment of sites corresponding to known motifs with stage-specific TREs (relative to a comparative stage). More specifically, we used function *findMotifsGenome*.*pl* (genome = hg38 and size = given) and the TREs which are shared or unique to the comparative stage were used as background.

### Transcription factor cistrome analysis

For a TF of interest, the binding motifs present in the stage-specific TREs were identified and the density of the motif was determined by the number of motifs within the window of +100 kb and -100 kb around the TSS for all the genes which are actively transcribed in the stage of interest (TPM > 50). The cistrome of a TF is defined using the following criteria: (1) genes have motif density > 0, (2) genes are actively transcribed (TPM > 50 in the ChRO-seq) and expressed (base mean > 500 in the RNA-seq) in the stage of interest, and (3) the genes are significantly uptranscribed and upregulated (padj < 0.2, p < 0.05, fold change > 1.5 in both ChRO-seq and RNA-seq by DESeq2) in the stage of interest relative to a comparison stage.

### Single cell RNA-seq data analysis

The scRNA-seq analysis of primary human fetal SI tissues was curated using recent, pre-existing datasets [13, 40, 41]. Briefly, the scRNA-seq dataset of human developing SI across different regions and multiple gestational time points were generated by 10x Genomics platform and mapped to hg19 using default alignment parameters provided by Cell Ranger. Seurat (v3.1) package [70] was applied to the downstream analysis. Cells with greater than 5% mitochondrial transcript proportion and fewer than 1000 unique detected genes were excluded. Ribosomal genes, mitochondrial genes, and genes located on sex chromosomes were also removed before data integration into a Seurat object. The cell type annotation of the dataset was pre-defined and detailed in [13]. In the present study, we focused on the data of matched human fetal duodenum and ileum samples (week 11, 14 and 19 of gestation).

### Statistics

All the padj and p-values presented in this study were determined by Wald test (DESeq2), unless otherwise specifically noted.

## Availability of data, materials and computation scripts

Raw and processed data generated in the sequencing studies will be made available upon publication.

## Author Contributions

Conceptualization, Y.-H.H., P.S.; Cell culture, S.H.; Cell sorting, M.K.D.; Chromatin isolation and library preparation for sequencing studies, Y.-H.H.; Bioinformatic analyses and data curation, Y.-H.H.; Resources and experiments of *Procr* reporter mice, Q.C.Y. and Y.A.Z.; Resources of scRNA-seq data, Q.Y., J.G.C. and J.R.S.; Writing (original draft), Y.-H.H., P.S.; Review and editing, Y.-H.H., J.R.S., and P.S.; Supervision, P.S.; Funding acquisition, Y.-H.H., J.R.S., and P.S.

## Acknowledgments

We would like to thank members of the Sethupathy laboratory, most notably Dr. Michael Shanahan, Dr. Ajeet Singh, and Dr. Matt Kanke, for helpful comments and feedback on the study and manuscript. We specially thank Dr. Tim Dinh in the Sethupathy laboratory for help in developing the bioinformatic analysis pipeline of ChRO-seq data and providing technical consultation on bioinformatics, Dr. Matt Kanke for providing additional technical consultation on bioinformatics, and Ramja Sritharan in the Sethupathy laboratory for offering technical consultation on the ChRO-seq protocol. Additionally, we are grateful to the Danko laboratory, specifically Ed Rice and Dr. Charles Danko, for helpful conversations on improving implementation of the ChRO-seq protocol. Finally, we also thank Dr. Jen Grenier and the Cornell Transcriptional Regulation & Expression Facility for RNA sequencing support; Michael Dellheim of the University of Michigan BRCF Flow Cytometry Core; Gina Newsome, Erika Katz, Maliha Berner, and Angeline Wu of the Michigan Medicine Translational Tissue Modeling Laboratory; and a University of Michigan funded initiative (Center for Gastrointestinal Research, Office of the Dean, Comprehensive Cancer Center, Departments of Pathology, Pharmacology, and Internal Medicine) with support by the Endowment for Basic Sciences.

---

**Supplementary Figure 1.**
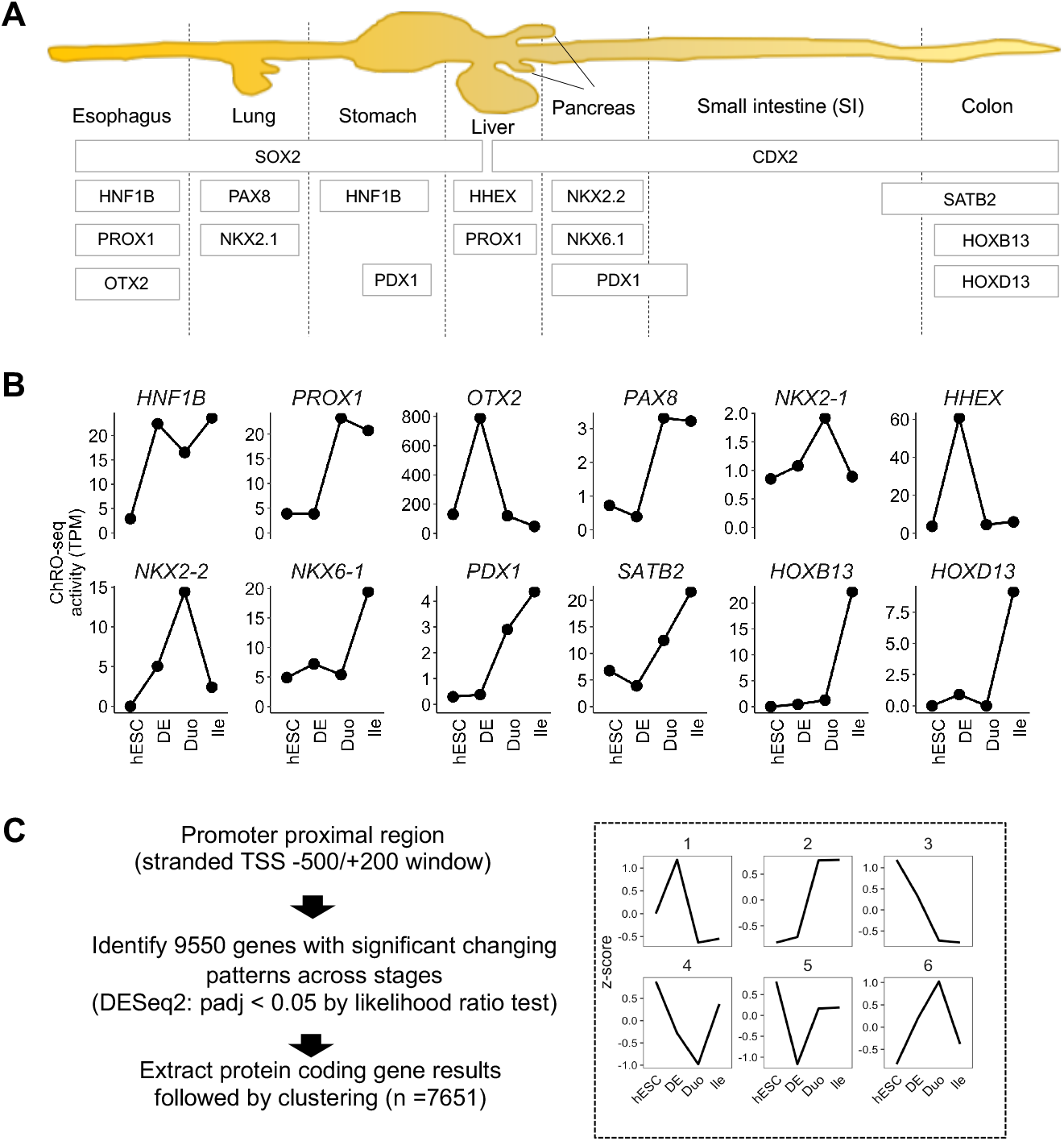
ChRO-seq profiling shows specificity of transcriptional markers in the directed differentiation model of human developing SI. (A) TFs associated with different gastrointestinal organoids that are generated by the hPSC directed differentiation methods^45-51^. (B) Transcriptional activity (ChRO-seq signal) of genes encoding the TFs shown in (A) across different stages. (C) Characterization of genes with distinct patterns of promoter activity across different stages based on the likelihood ratio test (DESeq2). The protein-coding subset of genes (n = 7,651) were clustered into different groups based on the changing patterns of promoter activity. ChRO-seq study: hESC, n = 3; DE, n = 4; Duo spheroid (Duo), n = 3; Ile spheroid (Ile), n = 3. RNA-seq study: hESC, n = 2; DE, n = 3; Duo, n = 6; Ile, n = 4. TPM, transcripts per million.

**Supplementary Figure 2.**
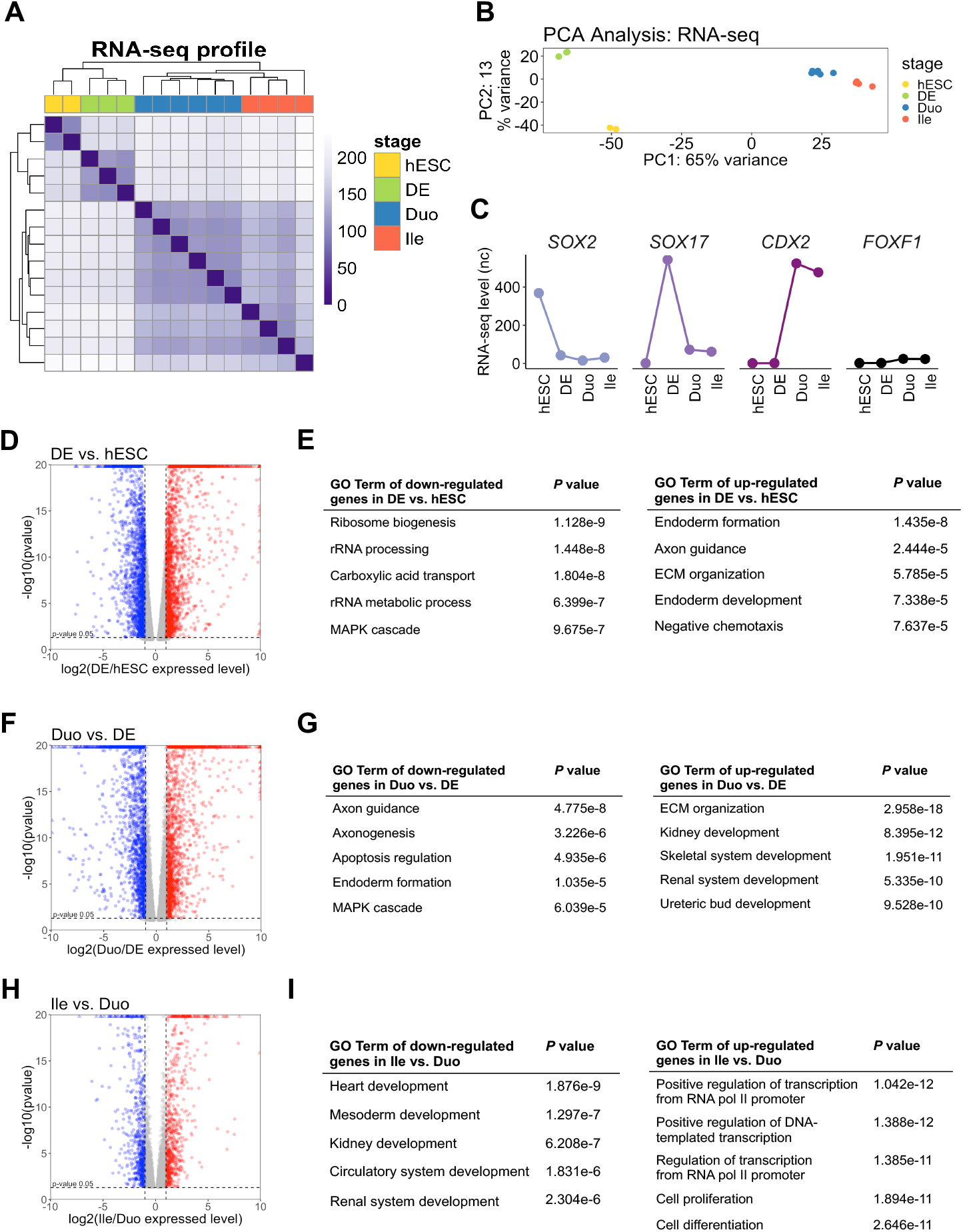
Steady-state gene expression profiles across all stages in the directed differentiation model of the developing human SI. (A) Hierarchical clustering analysis of RNA-seq expression profiles across stages. Color shade denotes sample to sample distance. (B) PCA of profiles of RNA-seq across stages. (C) RNA-seq expression levels of *SOX2, GATA6, SOX17, CDX2* and *FOXF1* across stages. (D, F, H) Volcano plot showing differentially expressed genes in the indicated comparison. Numbers in red and blue are number of up- and down-regulated genes in the comparison (base mean > 100, log_2_ fold change of transcription > 1, padj < 0.2 and p < 0.05 by Wald test; DESeq2). (E, G, I) Pathway enrichment analyses of up- and down-regulated genes in the indicated comparisons (GO term = GO Biological Process 2018). hESC, n= 2; DE, n= 3; Duo spheroid (Duo), n = 6; Ile spheroid (Ile), n = 4. nc, normalized counts.

**Supplementary Figure 3.**
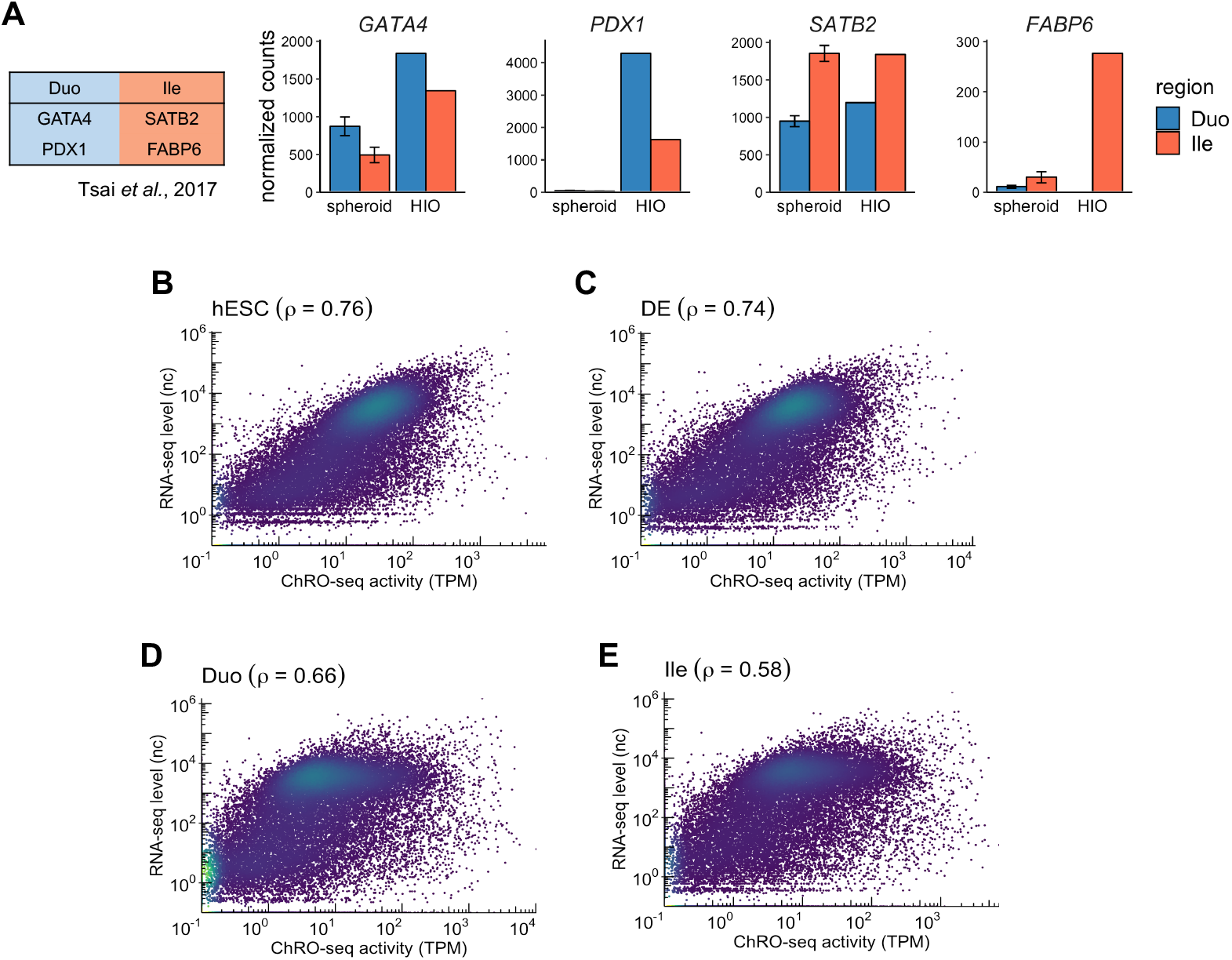
Integrative analysis of ChRO-seq and RNA-seq defines marker genes in the directed differentiation model of the developing human SI. (A) RNA-seq expression of genes associated with SI regional identity in spheroids and in HIOs with either Duo or Ile identity. Genes associated with SI regional identity in primary human fetal gut as well as in transplanted HIOs in Tsai *et al*., 2017^21^. Duo and Ile HIOs were generated by culturing Duo and Ile spheroids in 3D Matrigel for 28 days and purified by EPCAM+ (epithelial marker) sorting. The HIO samples were from the same batch used for ChRO-seq and RNA-seq of Duo and Ile spheroids in this study. (B-E) Genome wide correlation of transcribed levels (ChRO-seq) and expressed levels (RNA-seq) in the indicated stages. No expression or transcription level thresholds were used for genes to be included in the analysis. ChRO-seq study: hESC, n= 3; DE, n= 4; Duo spheroid (Duo), n = 3; Ile spheroid (Ile), n = 3. RNA-seq study: hESC, n= 2; DE, n= 3; Duo, n = 6; Ile, n = 4; Duo HIO, n = 1; Ile HIO, n = 1.

**Supplementary Figure 4.**
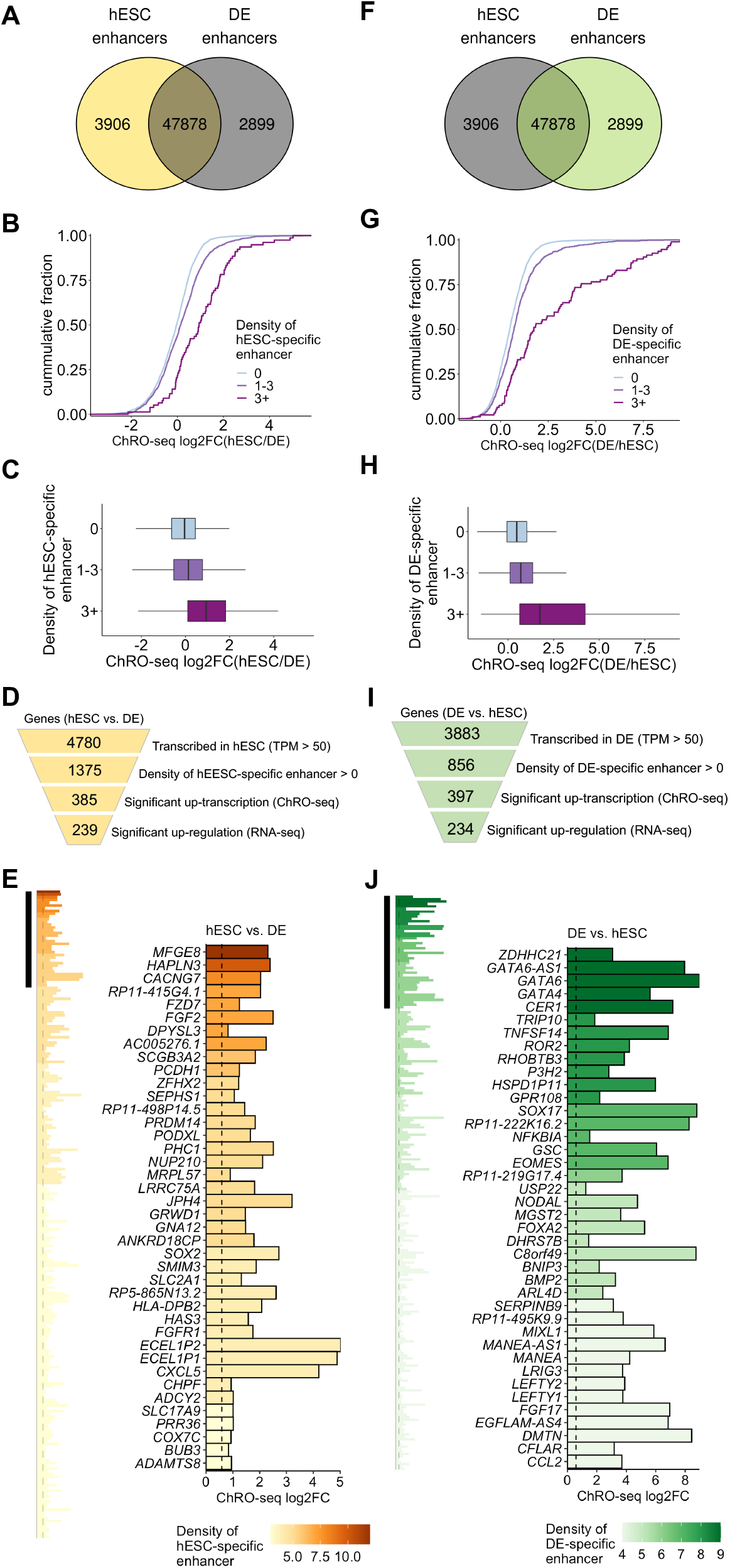
Identification of genes associated with stage-specific enhancers relevant to the stem cell and definitive endoderm stages. (A) Venn diagram showing stage-specific and shared enhancers between hESC and DE. hESC-specific enhancers (n = 3906) were defined in the comparison with DE. (B) Cumulative distribution of ChRO-seq fold change in the transcriptional activity of genes grouped into three different categories of hESC-specific enhancer density in hESC vs. DE. (C) Boxplot of ChRO-seq fold change in the transcriptional activity of genes grouped into three different categories of hESC-specific enhancer density in hESC vs. DE. (D) Identification of genes associated with hESC-specific enhancers (n = 239). (E) Bar graph showing genes associated with hESC-specific enhancers (left panel). Top 30 genes based on enhancer density are highlighted (right panel). (F) Venn diagram showing stage-specific and shared enhancers between hESC and DE. DE-specific enhancers (n = 2899) were defined in the comparison with hESC. (G) Cumulative distribution of ChRO-seq fold change in transcriptional activity of genes grouped into three different categories of DE-specific enhancer density in DE vs. hESC. (H) Boxplot of ChRO-seq fold change in transcriptional activity of genes grouped into three different categories of DE-specific enhancer density in DE vs. hESC. (I) Identification of genes associated with DE-specific enhancers (n = 234). (J) Bar graph showing genes associated with DE-specific enhancers (left panel). Top 30 genes based on enhancer density are highlighted (right panel). ChRO-seq study: hESC, n= 3; DE, n= 4. RNA-seq study: hESC, n= 2; DE, n= 3.

**Supplementary Figure 5.**
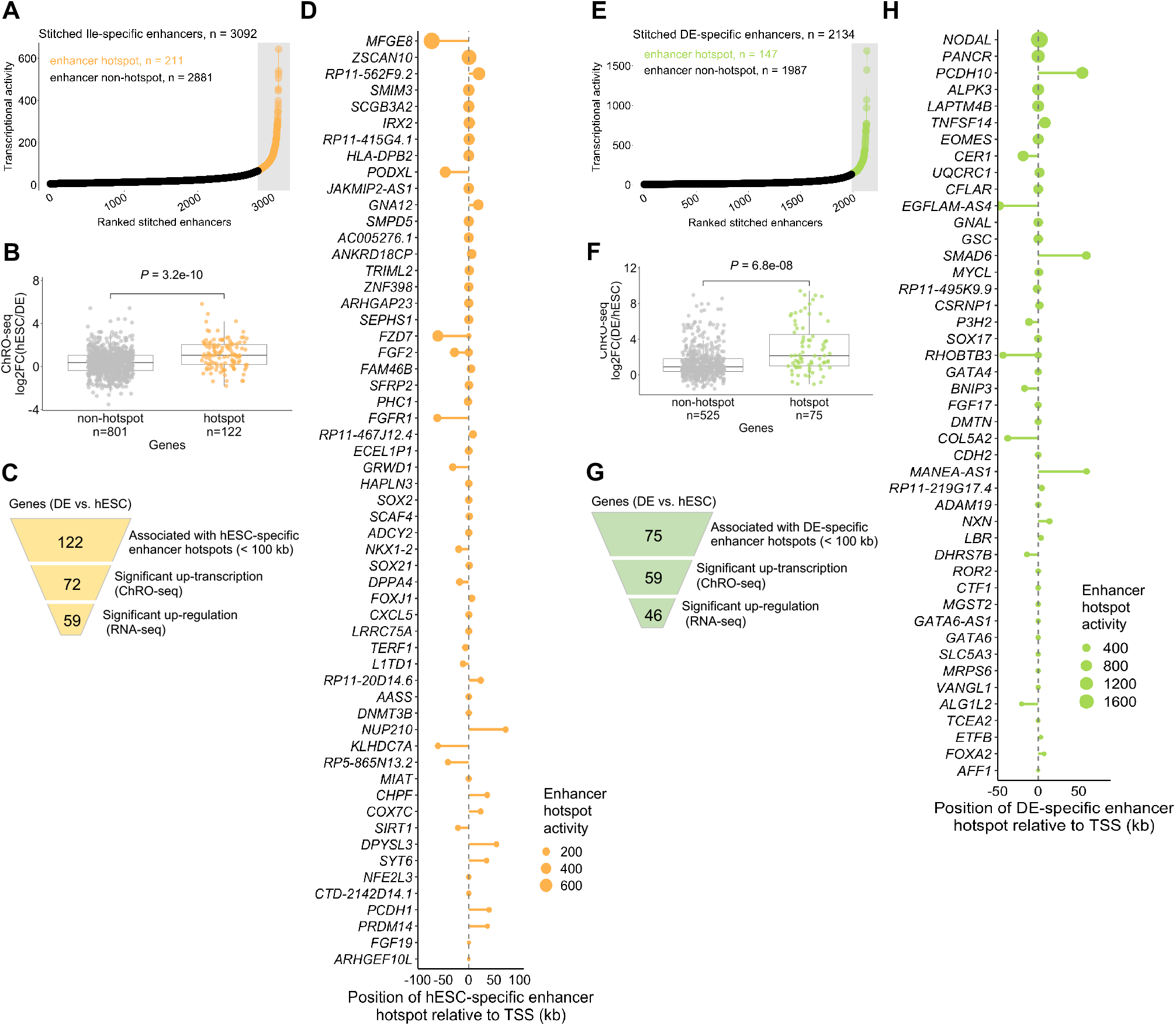
Identification of hESC- and DE-specific enhancer hotspots and associated genes. (A) hESC-specific stitched enhancers are ranked by transcriptional activity (ChRO-seq signal). The stitched enhancers with the highest transcription activity are defined as hESC-specific enhancer hotspots (n = 211; yellow) and the rest are enhancers non-hotspots (n = 2881; black). (B) ChRO-seq fold change in the transcriptional activity of genes associated with hESC-specific stitched enhancers, non-hotspots vs. hotspots (Wilcoxon test). (C) Identification of genes associated with hESC-specific enhancer hotspots. (D) Genes associated with hESC-specific enhancer hotspots (n=59). Relative position between enhancer hotspots and TSSs of the associated genes are shown. Dot size denotes transcriptional activity of a given DE-specific enhancer hotspot. (E) DE-specific stitched enhancers are ranked by transcriptional activity (ChRO-seq signal). The stitched enhancers with the highest transcription activity are defined as DE-specific enhancer hotspots (n = 147; green) and the rest are enhancer non-hotspots (n = 1987; black). (F) ChRO-seq fold change in the transcriptional activity of genes associated with DE-specific stitched enhancers, non-hotspots vs. hotspots (Wilcoxon test). (G) Identification of genes associated with DE-specific enhancer hotspots. (H) Genes associated with Ile-specific enhancer hotspots (n=46). Relative position between enhancer hotspots and TSSs of the associated genes are shown. Dot size denotes transcriptional activity of a given DE-specific enhancer hotspot. ChRO-seq study: hESC, n= 2; DE, n= 4. RNA-seq study: hESC, n=2; DE, n= 3.

**Supplementary Figure 6.**
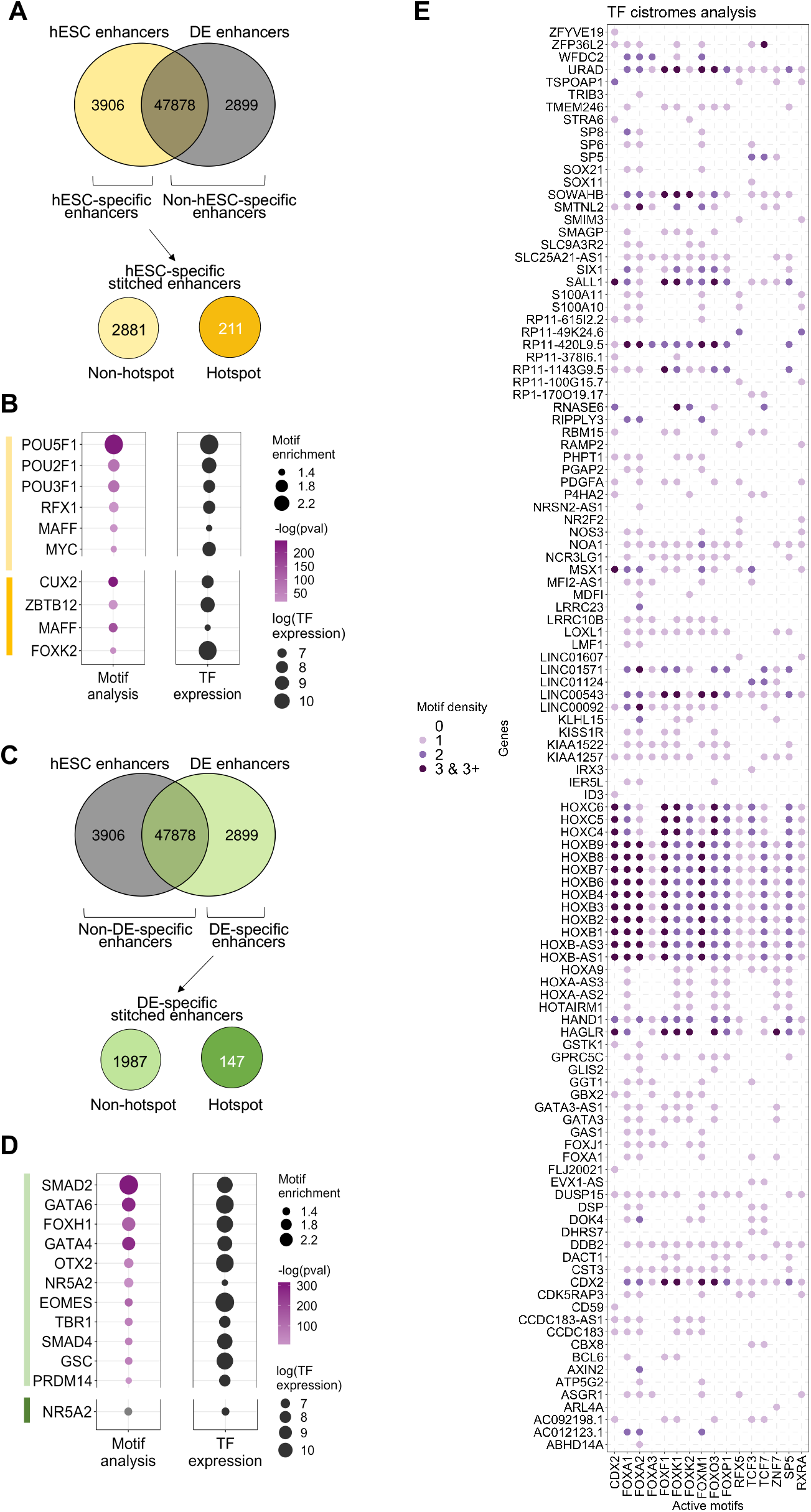
TF binding motif enrichment analysis of TREs specific to hESC or DE. (A) TF motif enrichment analyses were performed in hESC-specific enhancers (n = 3906) relative to non-hESC-specific TREs (n = 50,777) as well as in hESC-specific enhancer hotspots (n = 211) relative to non-hotspots (n = 2,881). (B) Motifs significantly enriched in hESC-specific enhancers (highlighted by light orange bar) or in hESC-specific enhancer hotspots (highlighted by dark orange bar). The steady-state expression (RNA-seq) of the corresponding TFs are also shown. (C) TF motif enrichment analyses were performed in DE-specific enhancers (n = 2899) relative to non-DE-specific TREs (n = 51,784) as well as in DE-specific enhancer hotspots (n = 147) relative to non-hotspots (n = 1987). (D) Motifs significantly enriched in DE-specific enhancers (highlighted by light green bar) or in DE-specific enhancer hotspots (highlighted by dark green bar). (E) Bubble plot showing full list of genes associated with active binding motifs of TFs that exhibit an overall enrichment of binding motifs in Duo-specific enhancers (defined in **Figure 6C**). ChRO-seq study: hESC, n= 3; DE, n = 4; Duo spheroid, n = 3.

